# Spatial multi-omics and deep learning reveal fingerprints of immunotherapy response and resistance in hepatocellular carcinoma

**DOI:** 10.1101/2025.06.11.656869

**Authors:** Zhenqin Wu, Joseph Boen, Sonali Jindal, Sreyashi Basu, Matthew Bieniosek, Siyu He, Michael LaPelusa, Aaron T. Mayer, Ahmed O. Kaseb, James Zou, Padmanee Sharma, Alexandro E. Trevino

## Abstract

Despite advances in immunotherapy treatment, nonresponse rates remain high, and mechanisms of resistance to checkpoint inhibition remain unclear. To address this gap, we performed spatial transcriptomic and proteomic profiling on human hepatocellular carcinoma tissues collected before and after immunotherapy. We developed an interpretable, multimodal deep learning framework to extract key cellular and molecular signatures from these data. Our graph neural network approach based on spatial proteomic inputs achieved outstanding performance (ROC-AUC > 0.9) in predicting patient treatment response. Key predictive features and associated spatial transcriptomic profiles revealed the multi-omic landscape of immunotherapy response and resistance. One such feature was an interface niche expressing restrictive extracellular matrix factors that physically separates tumor tissue and lymphoid aggregates in nonresponders. We integrate this and other spatially-resolved signatures into SPARC, a multi-omic “fingerprint” comprising scores for immunotherapy response and resistance mechanisms. This study lays groundwork for future patient stratification and treatment strategies in cancer immunotherapy.

## Introduction

Hepatocellular carcinoma (HCC) is the most prevalent type of liver cancer. Incidence of HCC has increased over the past several decades, and despite advances in treatment, including more inclusive surgery options and immunotherapies, mortality remains high, with a five-year survival rate of 18%[1,2]. The pathological and molecular presentation of HCC is variable, with clinicians recognizing at least 5 histological subtypes[3]. HCC cases can be further divided according to etiology[4,5], with up to 90% of patients having accompanying liver cirrhosis due to infection[6], metabolic disease[7], or other factors.

Immunotherapy has proved promising for both early and late stage HCC[8–10], though a significant fraction of HCC patients undergoing treatment fail to respond[1,11,12]. Understanding the cellular and molecular dynamics at play during therapy could lead to fewer failed outcomes, more personalized treatments, and mechanistic insight into disease processes. Previous studies have identified associations between immunotherapy response and diverse biomarkers[13]—including mutation burden[14,15], molecular signatures[16–18], immune cell states[19–22], pathological features[23], transcriptomics[24–27], and spatial organization[28–32]—in HCC and other cancers. Still, no validated clinical biomarker currently exists to guide precision medicine, and therapy-resistant patients have few options.

To explore the biology of immunotherapy further, we performed spatially-resolved, multi-omic profiling on tissue samples from HCC patients undergoing perioperative checkpoint inhibition[33,34]. We aligned and integrated data from spot-based spatial transcriptomics[35] (ST, 10X Visium), multiplexed immunofluorescence (MIF)-based spatial proteomics[36] (Akoya Phenocyler), and histopathology performed on consecutive adjacent sections. To analyze these data, we modeled treatment responses using interpretable graph neural networks and multilayer perceptrons, which mapped local cell microenvironments to predictions for response. We then classified these microenvironments, characterized their cellular organization and gene expression profiles, and assessed their relevance to immunotherapy response. Finally, we defined multifaceted cellular and transcriptomic signatures of HCC using these niches. These signatures span key biological processes and structural features—including immune cell infiltration and activation, lymphoid aggregate organization, extracellular matrix (ECM) remodeling, metabolism, and angiogenesis—that correlate with treatment response. We further revealed the dynamics of these signatures during the course of successful and unsuccessful immunotherapy. Our results suggest new approaches for patient stratification and therapy development in HCC.

## Results

The results are organized as follows: We first describe the spatially-resolved, multi-omic profiling of HCC immunotherapy recipients, highlighting histological, proteomic, and transcriptomic characteristics (**Fig. 1**). Next, we identify cell types from MIF data and compare them with co-registered ST data to assess alignment between modalities (**Fig. 2**). With these fundamentals established, we then employ predictive modeling to uncover neighborhood-scale determinants of response and resistance, termed microenvironment archetypes (**Fig. 3**). We identify and interpret key tumor- and lymphoid aggregate-associated microenvironment archetypes (**Figs. 4-5**), revealing response/resistance-associated cellular, proteomic, and transcriptomic signatures. Finally, we integrate signatures into a comprehensive fingerprint that we call SPARC to characterize distinct initial conditions and treatment-induced dynamics in HCC patients undergoing immunotherapy (**Fig. 6**).

**Figure 1:**
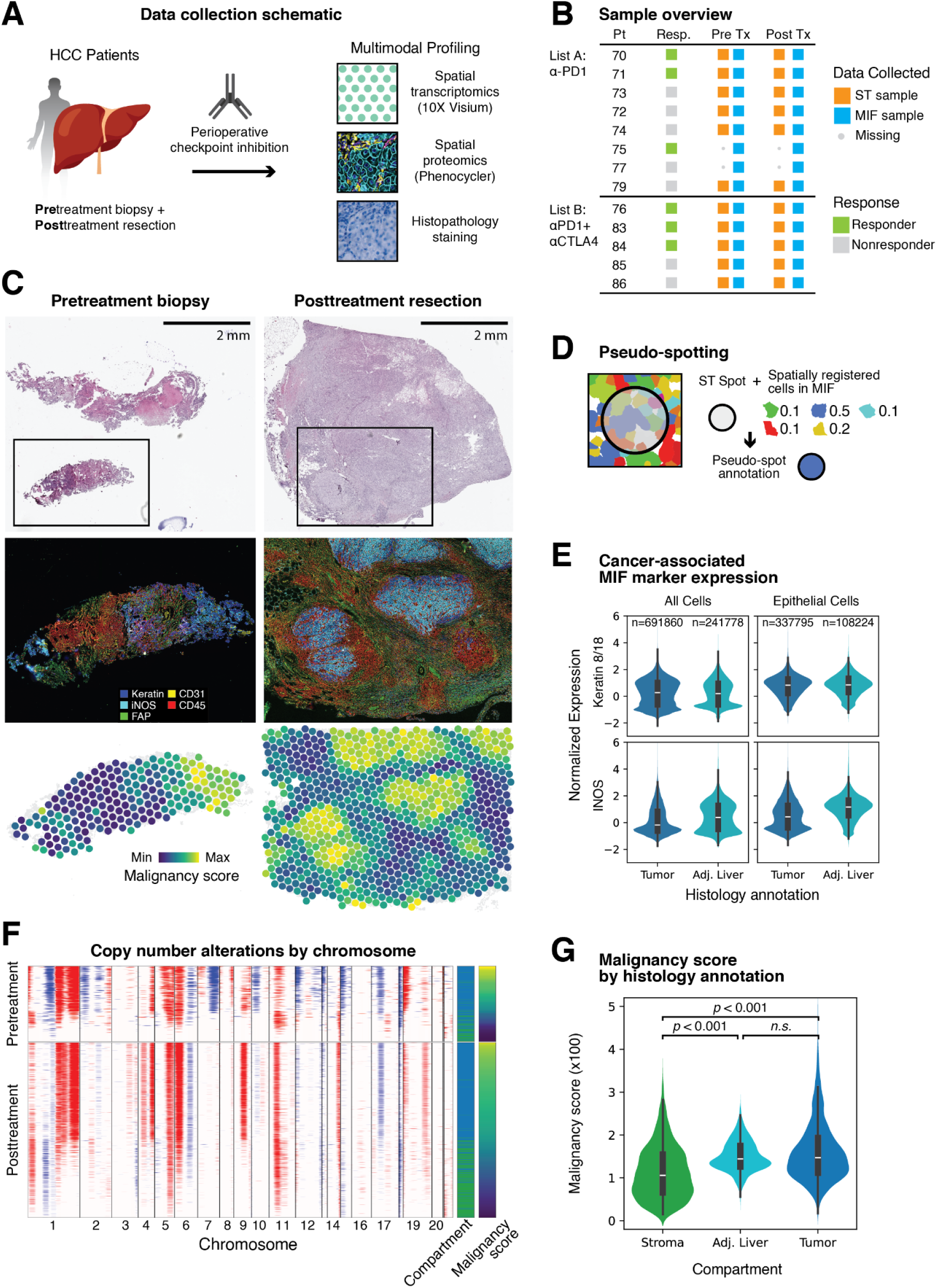
Pathological and clinical features of HCC patients before and after perioperative immunotherapy. **A.** Schematic of patient treatment and data collection. **B.** Sample overview including treatment (anti-PD1, anti-PD1 + anti-CTLA4), response status, and omics modalities collected. **C.** Example pathology stains (top), MIF stains (middle), and spatially co-registered ST spots (bottom) of a pretreatment biopsy and a posttreatment resection from the same patient. Major protein biomarkers and per-spot malignancy scores (**Methods**) are visualized. **D.** Schematic of pseudo-spotting: ST spots were annotated based on the major cellular annotation class derived from spatially overlapping cells identified in MIF. **E.** Heterogeneous expression of cancer-associated markers in epithelium. Keratin is consistently expressed in epithelial cells. INOS expression is observed in the epithelium of tumor-adjacent liver, as well as in a subset of tumoral epithelial cells. **F.** Copy number alteration analysis of ST samples from the representative patient revealing amplification (red) and deletion (blue) signals. Rows represent ST spots, and columns are genes ordered by chromosomal coordinates. Malignancy scores and histology annotations of spots are visualized as side columns. **G.** Summary of malignancy scores by histology annotations. Malignancy scores are significantly higher in both tumor and tumor-adjacent liver compartments (MWU test, *p*<10^-3^).

**Figure 2:**
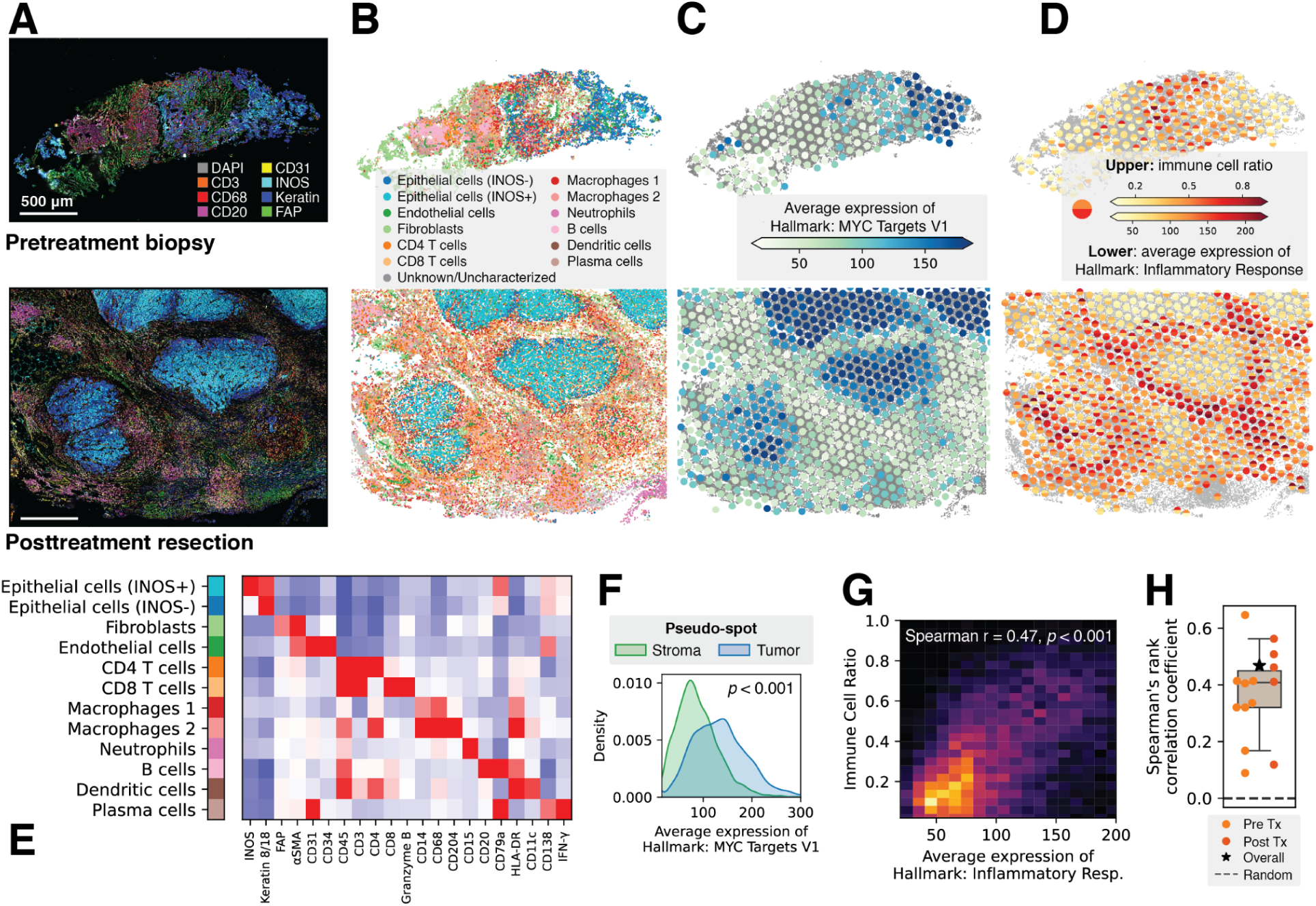
Multimodal cellular and molecular characterization of HCC. **A.** MIF images of a pretreatment biopsy and a posttreatment resection sample from the same patient. Key biomarkers including Keratin (blue), INOS (cyan), CD31 (yellow), FAP (green) and various immune cell markers are displayed. **B.** Scatter plots showing spatial distributions of cells in the representative samples. Cells were segmented and annotated based on integrated protein expression from MIF. **C.** Co-registered ST spots plotted over the cell annotations. Each spot is colored by the aggregated expression of genes in a tumoral hallmark gene set: MYC targets V1. Note the spatial alignment between epithelial cells and higher-expressing spots. **D.** The same set of ST spots colored by ratios of immune cells in pseudo-spots (top) and aggregated expression of a hallmark gene set: inflammatory response (bottom). The two quantities exhibit high spatial concordance. **E.** Heatmap showing signature protein expression from diverse cell types. **F.** Kernel density plots showing expression of tumoral hallmark genes in ST spots. Spots were classified based on the identities of their spatially co-registered cells (i.e., pseudo-spotting). Tumoral spots exhibit significantly higher expression (MWU test, *p* < 10^-3^). A similar comparison conducted on spots annotated by histology compartments suggested consistent differences. **G.** Heatmap showing the associations between immune cell ratios and immune gene signatures. For each spot, the ratio of immune cells in its pseudo-spot and gene expression in an immune-focused hallmark gene set were calculated. The two quantities show high concordance, with a significant positive correlation (Spearman’s rank correlation coefficient 0.47, *p* < 10^-3^). **H.** Box plot showing consistent associations between immune cell ratios and immune gene signatures across samples.

**Figure 3:**
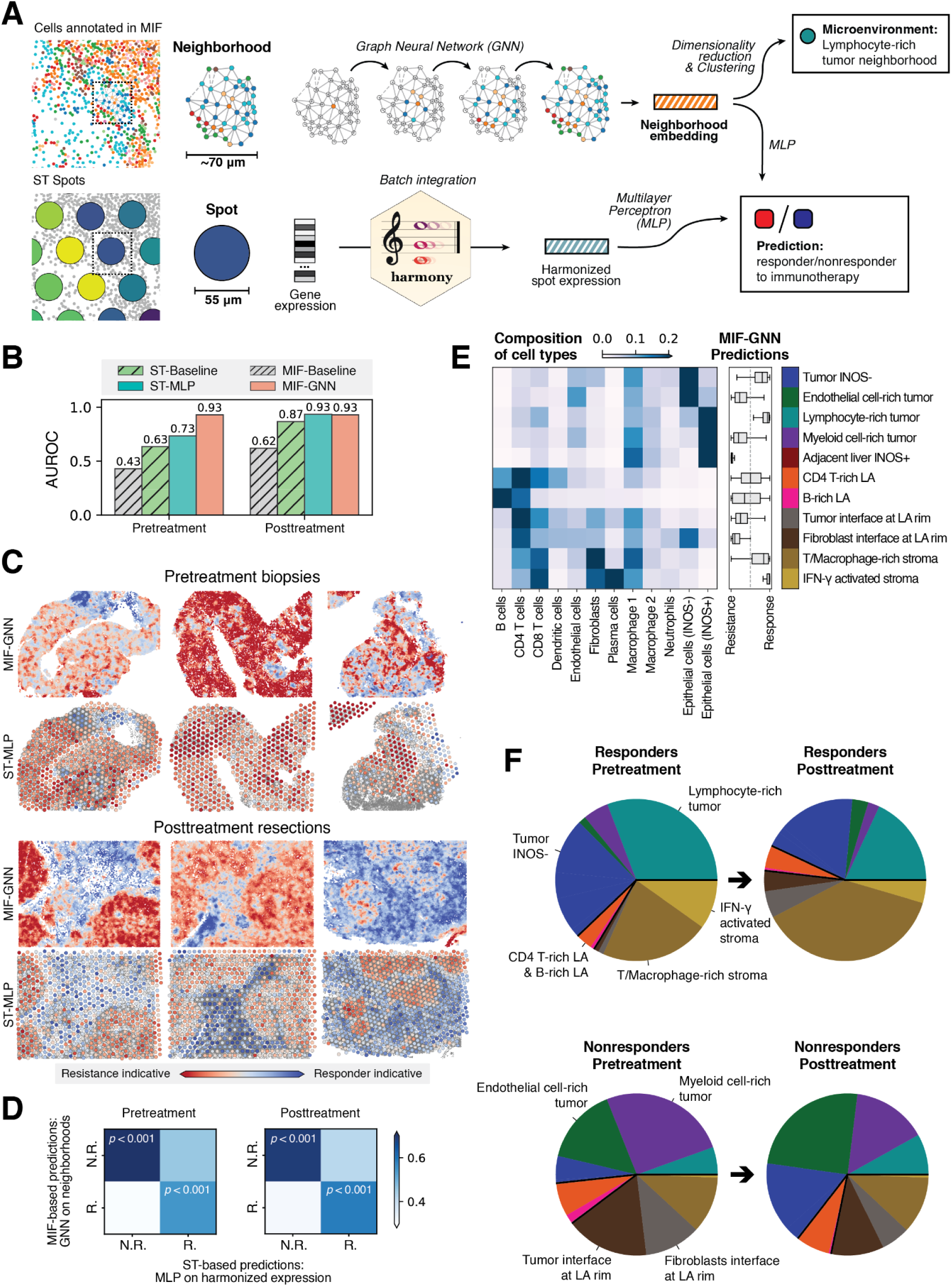
Multi-modal predictive modeling of responses to immunotherapy. **A.** Schematic of multi-modal predictive modeling. Spatial neighborhoods of cells characterized in MIF were comparable in spatial dimensions to ST spots. They were extracted and processed by modality-specific deep learning models to predict responses to immunotherapy. Microenvironment archetypes were defined by clustering embeddings of neighborhoods. **B.** Bar plots showing performances from sample-scale (i.e., baselines) and neighborhood-scale predictors. MIF-GNN achieved the best AUROC in both conditions. **C.** Predictions on representative regions visualized as scatter plots. Note the spatial concordances between predictions from the two modalities. **D.** Heatmap showing concordances between neighborhood-based and spot-based predictions. Neighborhood-scale predictions of immunotherapy susceptibility were aggregated across all pretreatment or posttreatment samples, exhibiting significant spatial concordance (t-test, *p* < 10^-3^). **E.** Heatmap showing cell type compositions of microenvironment archetypes, defined by clustering MIF-GNN embeddings. Box plots in the middle show predictions for response/resistance from different microenvironment archetypes. **F.** Pie charts showing the differences and dynamics in microenvironment archetype compositions across conditions.

**Figure 4:**
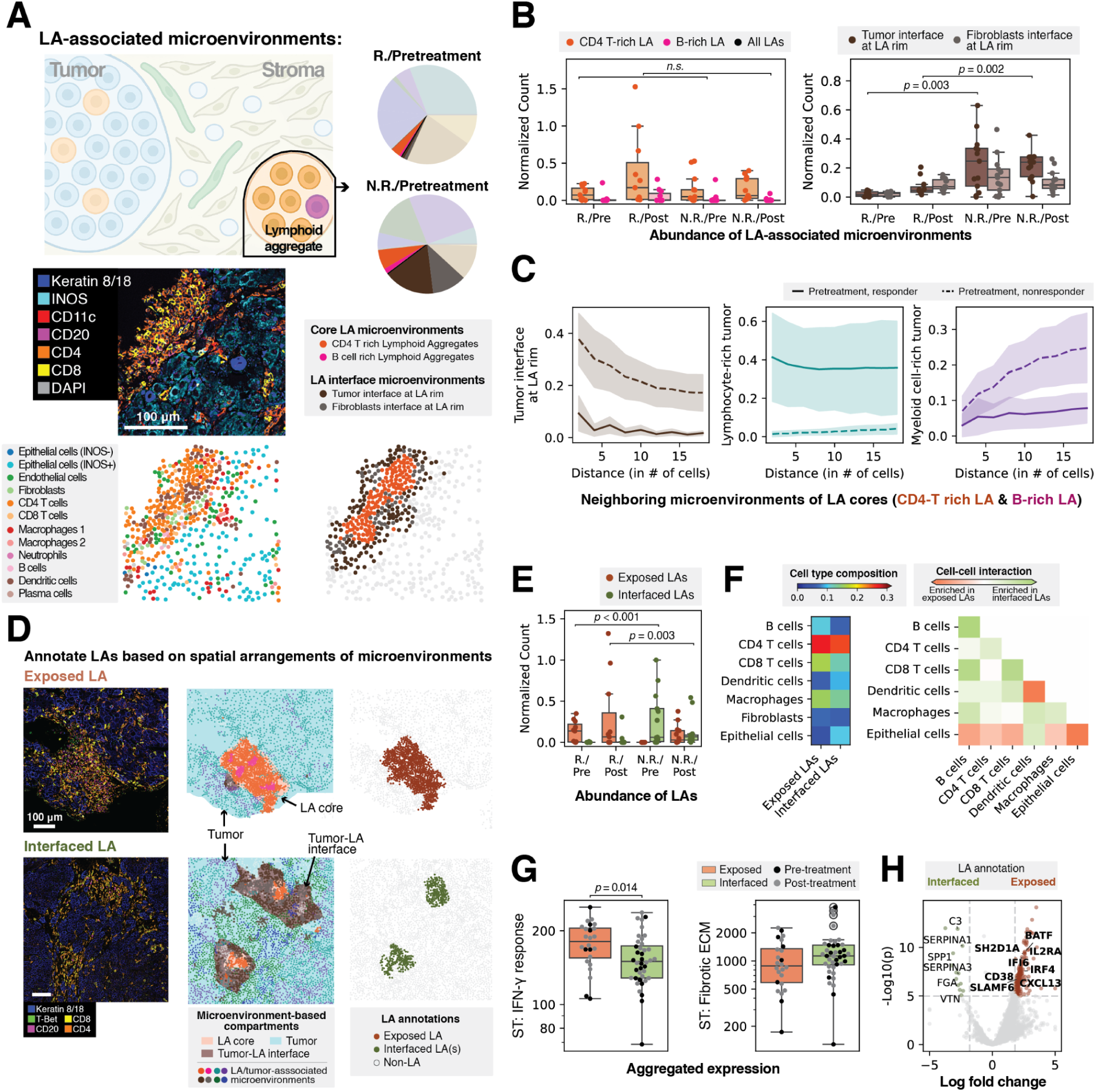
Structures of lymphoid aggregates (LAs) are associated with T cell activities and resistance to immunotherapy. **A.** Schematic of LA-associated microenvironment archetypes, accompanied by an example LA with cell type and microenvironment annotations. Two archetypes were identified in the core of LAs, while the other two appeared on the periphery. **B.** Abundance of LA-associated microenvironment archetypes. CD4 T-rich and B-rich LA show no significant enrichment. Tumor interface microenvironments were significantly enriched in nonresponders (MWU test, pretreatment: *p* = 0.003, posttreatment: *p* = 0.002). **C.** Different spatial arrangements of microenvironments around LAs. Compositions of microenvironment archetypes in the periphery of LA cores (CD 4 T-rich LA and B-rich LA) at different distances were visualized. Tumor interface microenvironments were present in the vicinity of LAs, especially in nonresponders. In contrast, LAs in responders were more frequently positioned adjacent to the tumor (i.e., lymphocyte-rich tumor microenvironments, detailed in the next section). **D.** Annotations of LAs based on spatial arrangements of microenvironments. LAs were classified as exposed or interfaced based on the presence of tumor-LA interfaces (i.e., brown area in the middle column). **E.** Abundance of exposed and interfaced LAs. Exposed LAs were more frequently observed in responders, whereas interfaced LAs were prevalent in nonresponders. **F.** Cell type compositions and cell-cell interaction patterns in exposed and interfaced LAs. Exposed LAs have more B cells and CD8 T cells, alongside stronger immune-tumor interactions. **G.** Gene expression in relevance to interferons. We identified significantly higher expression of IFN-γ response related genes in exposed LAs (MWU test, pretreatment: *p* = 0.014). Interfaced LAs showed higher expression of fibrotic ECM genes. **H.** Differential gene expression analysis between exposed and interfaced LAs. Genes enriched in exposed LAs are annotated in bold.

**Figure 5:**
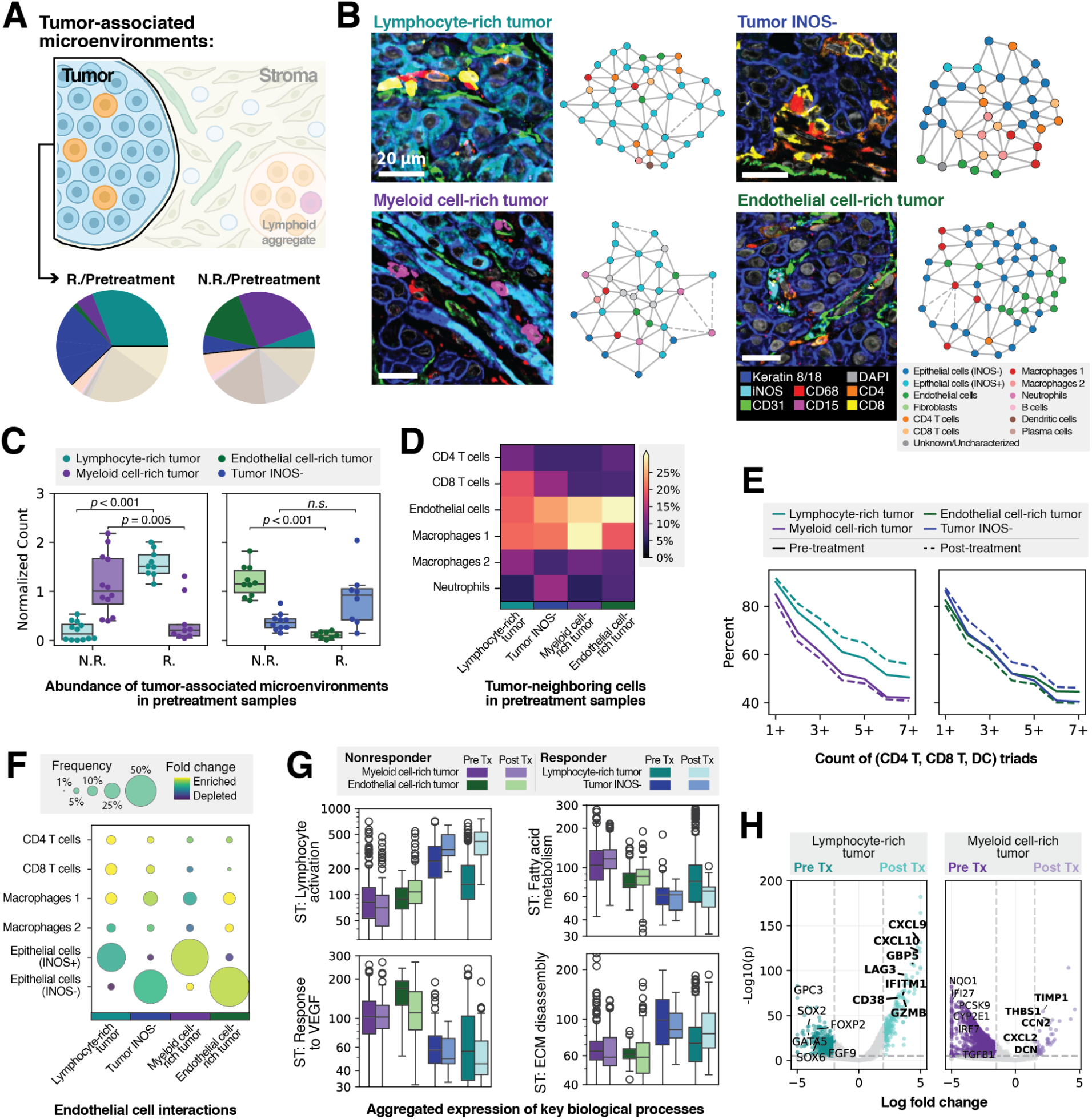
Tumor-associated microenvironments with distinct immune environments and gene signatures. **A.** Schematic of tumor-associated microenvironment archetypes. **B.** Examples of the four tumor-associated microenvironment archetypes. **C.** Abundance of tumor-associated microenvironments in pretreatment samples. Lymphocyte-rich tumor microenvironments were significantly enriched in responders (MWU test, *p* < 10^-3^). Myeloid cell-rich and endothelial cell-rich tumor microenvironments were significantly enriched in nonresponders (MWU test, *p* = 0.005 and *p* < 10^-3^). **D.** Compositions of tumor-neighboring cells in tumor-associated microenvironments. **E.** Abundance of immune triads (CD4 T cells, CD8 T cells, and dendritic cells) in tumor-associated microenvironments. Increases in immune triads after treatment were observed in response-associated microenvironments (lymphocyte-rich tumor and tumor INOS-). The reverse trend was observed in the resistance-associated microenvironments. **F.** Interactions of endothelial cells. Sizes of the circles indicate the frequency, whereas colors represent enrichment normalized by average composition. Endothelial cells predominantly interact with tumor cells in endothelial cell-rich and myeloid cell-rich tumor microenvironments. They interact more frequently with immune cells, especially T cells, in lymphocyte-rich tumor microenvironments. **G.** Aggregated expression of genes in key biological processes across tumor-associated microenvironments. **H.** Treatment-induced changes identified by differential gene expression analysis between pretreatment and posttreatment microenvironments. Posttreatment-enriched genes were annotated in bold.

**Figure 6:**
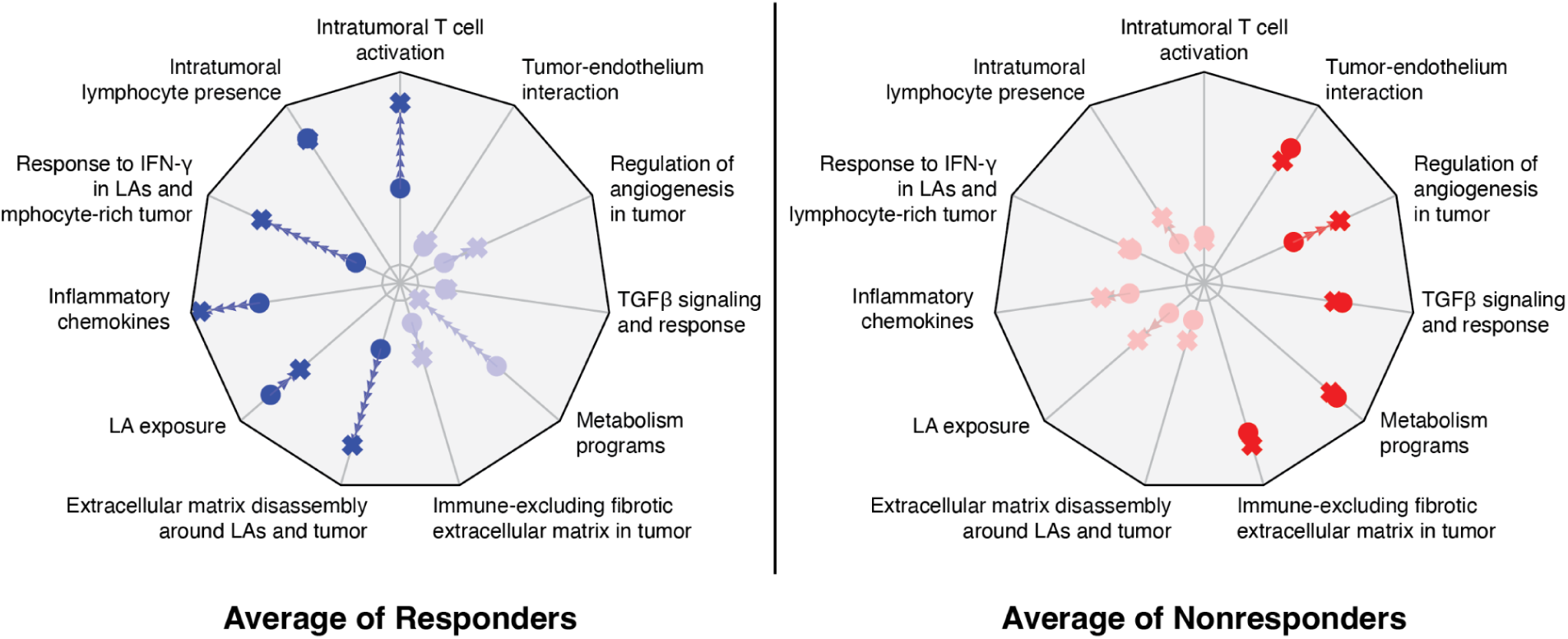
Fingerprints of response and resistance signatures. Radial plots summarizing key immune and tumoral signatures derived from microenvironment-based analysis. Each axis represents a distinct feature based on either MIF or ST inputs. The graph in blue represents responders, and the graph in red represents nonresponders. For each signature, circles indicate the pretreatment state and crosses indicate the posttreatment state, with arrows showing treatment-induced changes.

### Spatially-resolved, multi-omic profiling of HCC responders and nonresponders

We obtained tissue samples from a recent Phase II clinical trial in which nivolumab (anti-PD1) and nivolumab plus ipilimumab (anti-CTLA4) were evaluated for safety, with secondary endpoints of progression-free survival and overall response rate [34]. Major pathological response (MPR)—defined as complete response (100% tumor necrosis) or greater than 70% tumor necrosis at 6 weeks—was identified in a post-hoc analysis as a key indicator of therapeutic success. Accordingly, we denote MPR-positive patients as “responders”. Of the 20 trial patients who underwent pretreatment biopsy, treatment, and posttreatment resection, 13 were submitted for spatially-resolved, multi-omic analysis in this study, comprising 6 responders and 7 nonresponders. We performed hematoxylin and eosin (H&E) staining, 51-plex immunofluorescence (MIF, Akoya Phenocycler [36]), and spot-based spatial transcriptomics (ST, 10X Visium [35]) on sequential histological sections from both pretreatment and posttreatment samples for each patient (**Fig. 1A-C**). A stringent multi-stage quality assessment of tissues and samples (**Supplementary Fig. 1, Methods**) excluded some samples from further analysis (n = 2 field of views for MIF and 2 cases for ST), resulting in a final dataset of 48 MIF fields of view (24 pre and 24 posttreatment) and 22 ST samples (11 pre and 11 posttreatment).

To enable integrative analysis, we developed a computational pipeline to spatially co-register these modalities (**Methods, Supplementary Fig. 1F**), aligning and mapping data from different platforms to a shared spatial coordinate system. Of the 48 MIF fields of view, 27 were registered to ST data from adjacent tissue sections (**Supplementary Table 1**). To account for the differing spatial resolutions of the two modalities, we implemented a “pseudo-spotting” approach (**Fig. 1D**), wherein ST spots are mapped to spatially overlapping cells identified in MIF images. While the contents of adjacent sections are not expected to be identical, we assume they are correlated, and validate this assumption throughout the subsequent analysis.

### Visual and pathological signatures of HCC

Pretreatment biopsies were substantially smaller in size than posttreatment resection samples (**Fig. 1C**). Tumor, tumor-adjacent liver, and lymphoid aggregates (LAs) were manually annotated by a pathologist on H&E slides and confirmed on MIF data. LAs did not consistently express a panel of tertiary lymphoid structure (TLS) markers[37], and no mature TLSs were observed in these samples (**Supplementary Fig. 2**). In MIF data, we noticed the heterogeneous expression of inducible nitrous oxide synthase (INOS) specifically in Keratin 8/18-positive epithelial cells, with no expression observed in immune cell populations. While INOS-negative epithelial regions usually reflected tumor-annotated histology, INOS-positive epithelial regions were found in both tumor and tumor-adjacent liver compartments[38] (**Fig. 1E**).

To understand how genomic alterations were distributed in the tissue samples, we applied a computational copy number alteration detection tool[39] to ST data (**Fig. 1F**, **Methods**) and summarized the degree of amplification and duplication on the spot level as a malignancy score (**Fig. 1C**, **G**). Although malignancy scores varied widely across patients, there were no systematic differences between responders and nonresponders, or between pretreatment and posttreatment samples (**Supplementary Fig. 3**). Tumor compartments had significantly greater malignancy scores than stromal compartments (Mann–Whitney U/MWU test, *p*<10^-3^). Spots within tumor-adjacent liver compartments had malignancy scores nearly as high as tumoral spots, suggesting that tumor-adjacent liver may be extensively transformed in some patients, despite appearing histologically normal (**Fig. 1G**).

### Multimodal cellular and molecular characterization of HCC

Paired MIF and ST data enabled us to characterize protein expression, cell types, and gene expression levels *in situ* (**Fig. 2A-D**), and to compare these features according to histological or phenotypic landmarks. We identified cell types from MIF using unsupervised clustering of single-cell protein expression profiles[40] (**Fig. 2E**, **Supplementary Fig. 4A-C**, **Methods**). While cell compositions varied between samples and compartments, most cell types were represented in most samples (**Supplementary Fig. 4D-E**). Global cell-cell interaction statistics were profiled by summarizing neighboring pairs of cells observed in all MIF samples (**Supplementary Fig. 4F**), highlighting an overall bias of myeloid cell infiltration into tumor and enriched CD4-B cell interactions.

To illustrate and further verify the co-registration of our data types, we correlated MIF-based cell annotations with ST-based gene expression in space. Using MIF cell types, we created pseudo-spot annotations for tumor versus stroma, then compared the expression of a tumor hallmark gene set (“MYC Targets”) between these annotations (**Fig. 2C**, **F**). Similarly, we correlated MIF-based immune cell composition with inflammatory gene set expression in ST spots (**Fig. 2D, G-H**). As expected, both comparisons suggested substantial associations between co-registered MIF-based cellular views and ST-based spot views.

### Modeling associations between neighborhoods and immunotherapy response

To uncover the molecular and cellular signatures associated with immunotherapy response, we conducted a predictive modeling study leveraging our multimodal characterization. The goal was to identify features strongly linked to treatment response or resistance and to elucidate their underlying molecular and cellular biology.

To establish connections between treatment response and spatially-resolved cellular and transcriptomic features, we applied parallel deep learning models to neighborhood-scale spatial units (**Fig. 3A**), enabling us to identify areas of interest at fine spatial resolution. With MIF data, we adopted a graph neural network-based approach[30] to encode MIF cellular neighborhoods to predict response, hereafter referred to as MIF-GNN (**Methods**). In brief, we modeled each cell together with its neighboring cells within 3 hops, corresponding to a diameter of approximately 50 to 100 µm—comparable to the size of a single ST spot. For ST data, we deployed a multilayer perceptron model to predict patient response using gene expression from ST spots, hereafter referred to as ST-MLP (**Methods**).

We conducted predictive modeling of immunotherapy response using pretreatment and posttreatment samples separately, following a four-fold patient cross-validation strategy (**Methods**). Apart from the neighborhood-scale models, we designed baseline models that utilized sample-scale inputs such as cell type compositions across whole MIF regions or pseudo-bulk gene expression of aggregated ST spots (**Supplementary Fig. 5A**, **Methods**), allowing us to evaluate the added value of finer spatial resolution and spatial information. An additional cross-treatment validation was conducted to assess the imputed benefits of additional anti-CTLA4 treatment (**Supplementary Fig. 6**).

Both neighborhood-scale predictors demonstrated superior performances (**Fig. 3B**, **Supplementary Fig. 5B, E**) in predicting immunotherapy response, with the MIF-GNN approach achieving the best area under the receiver operating characteristic curve (AUROC): 0.93 in both pretreatment and posttreatment conditions. While sample-scale predictors failed to distinguish responders from nonresponders, MIF-GNN and ST-MLP approaches were highly capable of differentiating these groups (**Supplementary Fig. 5A, C, D, F**). Joint modeling of MIF and ST inputs was attempted, but the added performance benefit was marginal, likely due to the saturating performance of the MIF-GNN approach.

Neighborhood-scale predictions varied across spatial locations. Notably, the directionalities of MIF-GNN predictions and ST-MLP predictions were significantly correlated across space (**Fig. 3C-D, Methods**). In other words, response- and resistance-associated neighborhoods exhibit both cellular and transcriptomic signatures that were independently captured by the respective MIF and ST predictors.

### Disease-relevant microenvironments revealed by embedding analysis

Informed by the spatial variability of model predictions, we hypothesized that subsets of neighborhoods could be drivers of response or resistance. We reasoned that by jointly interpreting the cellular and transcriptional states of these key driver neighborhoods, we could gain insights into the underlying mechanisms of response or resistance. To achieve this, we clustered the MIF-GNN embeddings derived from the model that predicts response in pretreatment condition[30] (**Supplementary Fig. 7A**). We termed the resulting clusters “microenvironment archetypes”, or “microenvironments” for short (**Methods**).

These microenvironment archetypes exhibited diverse cell type compositions (**Fig. 3E**) and spatial organization patterns (**Supplementary Fig. 7B**), and we labeled them according to these factors. Our analysis includes four tumor-associated microenvironments (Tumor INOS-, Endothelial cell-rich tumor, Lymphocyte cell-rich tumor, and Myeloid cell-rich tumor); four immune-rich microenvironments (CD4 T-rich LA, B cell-rich LA, Tumor interface at LA rim, and Fibroblast interface at LA rim); two stromal microenvironments (T/Macrophage-rich stroma and INF-γ activated stroma); and an Adjacent liver INOS+ microenvironment. As expected, some microenvironments were consistently associated with response or resistance by the MIF-GNN model, while others were more neutral (**Fig. 3E**). Response or resistance-associated microenvironments were also enriched in specific patient groups–pre or posttreatment, responder or nonresponder–in terms of their prevalence (**Supplementary Fig. 8**). Overall, we observed markedly different distributions of microenvironment archetypes between responders and non-responders (**Fig. 3F**).

Through pseudo-spotting, we next curated ST spots that predominantly mapped to one microenvironment archetype (**Methods**). To reveal their associated gene expression signatures, we conducted both one-versus-rest comparisons (**Supplementary Fig. 9**, **Methods**) and targeted differential gene expression analyses, which will be detailed in the following sections.

### Interface structures in lymphoid aggregates restrict lymphocyte activation and infiltration during immunotherapy

LAs—organized clusters of immune cells, primarily B and T lymphocytes—play important roles in regulating immune activities within the tumor microenvironment[41], especially in the context of immunotherapies[42–45]. Among all microenvironment archetypes, we identified four that are relevant to LAs **(Fig. 4A**) and classified them according to their locations (**Supplementary Fig. 10A-C**) and cell type compositions (**Fig. 3E**, **Supplementary Fig. 7B**).

Neither the general abundance of LAs nor differences in their cell compositions were significantly linked to response (**Fig. 4B**). However, the presence of a specific tumor-LA interface microenvironment archetype was strongly associated with resistance (brown in **Fig. 4B**). Analysis of the spatial arrangement of LAs (**Fig. 4C**) revealed that nonresponder LAs had more tumor interface microenvironments in their vicinity, while responder LAs were often directly adjacent to tumor. Responder LAs were especially adjacent to a class of lymphocyte-rich tumor microenvironment detailed in the next section.

Given these observations about the tissue structure of LAs, we next annotated each individual LA as “exposed” or “interfaced” based on the presence or absence of a tumor-LA interface microenvironment (**Fig. 4D**, **Methods**). This new definition showed that exposed LAs were significantly more abundant in responders, while interfaced LAs were predominantly present in nonresponders (**Fig. 4E**).

To understand the distinctions between exposed and interfaced LAs further, we next analyzed the spatial organization of cells within these niches. The two types of LAs contained comparable amounts of CD4 T cells, but exposed LAs exhibited somewhat greater fractions of CD8 T cells (**Fig. 4F**), as well as higher *CD8A* gene expression (**Supplementary Fig. 11**). Direct immune-tumor interactions were more abundant in exposed LAs (**Fig. 4F**). *In silico* permutation followed by MIF-GNN model evaluation (**Methods**) supported these findings. For instance, synthetic mixing of tumor cells and lymphocytes (**Supplementary Fig. 12A-C**) or insertion of additional CD8 T cells (**Supplementary Fig. 12D-F**) mitigated associations with resistance.

To further assess LA functional states, we extracted and compared their protein and gene expression profiles (**Supplementary Fig. 11, Supplementary Fig. 10D-E**). While *CD4* and *CD3* expression were comparable across LAs, exposed LAs exhibited significantly higher expression of Th1- and Tfh-associated markers including *T-bet*/*TBX21* (green in **Fig. 4D**), *ICOS*, *STAT4*, *BCL6*, *CXCR5*, and *CXCL13.* Moreover, gene signatures relevant IFN-γ response were significantly higher in exposed LAs (**Fig. 4G** left, MWU test, *p* = 0.014). These findings align with previous reports of a *CD4*(+)*ICOS*(hi) T cell population during immune checkpoint blockade treatment[46–48]. In addition to strong INF-γ-related expression in exposed LAs, we identified an IFN-γ activated stromal microenvironment archetype that was observed exclusively in responders (**Fig. 3F**, **Supplementary Fig. 13A-B**). This archetype was not classified as an LA by histology or our model, but exhibited enrichment for immune cells, strong cellular interactions between CD8 T cells and IFN-γ positive immune cells (**Supplementary Fig. 13C**), and elevated IFN-γ signaling and T cell activation signatures by gene expression (**Supplementary Fig. 13D**). Collectively, these measurements show that exposed LAs are functionally distinct from interfaced LAs, and identify IFN-γ signaling as a key differentiating feature.

Results from differential expression analysis between exposed and interfaced LAs were consistent with these findings (**Fig. 4H**). Exposed LAs showed upregulation of genes related to T cell activation (e.g., *IL2RA*, *CD38*, *IFI6*) and Tfh function (e.g., *BATF*, *IRF4*, *SH2D1A*, *SLAMF6*), along with markers of tertiary lymphoid structure formation (e.g., *CXCL13*). In contrast, interfaced LAs exhibited fewer immune activation genes and showed elevated expression of ECM stabilization genes (e.g., *SERPINA1, SERPINA3, VTN, ITIH2*) and a macrophage-associated marker *SPP1*, which has been strongly implicated in tumor–immune barrier formation[21]. Further quantification indicated heightened expression of genes related to fibrotic ECM (**Methods**, **Supplementary Table 3**) in interfaced LAs (**Fig. 4G** right, **Supplementary Fig. 10E**). These findings suggest that the physical segregation of lymphocytes and tumor cells observed in interfaced LAs may be driven by the various ECM factors, which contributed to the establishment and maintenance of a protective barrier that insulates tumors from immune infiltration.

### Cellular and transcriptomic signatures in tumor-associated microenvironments indicate immunotherapy susceptibility

We next investigated microenvironment archetypes with substantial tumor cell fractions and explored their interplay with distinct types of LAs (**Fig. 5A**). Four tumor-associated microenvironment archetypes were identified: two in INOS-positive compartments (**Fig. 5B, Supplementary Fig. 14**) and the two in INOS-negative compartments (**Fig. 5B, Supplementary Fig. 15**), where INOS positivity did not strongly correlate with treatment response. A separate archetype was identified in tumor-adjacent liver tissue and exhibited unique spatial features (**Supplementary Fig. 17**).

Among these archetypes, we observed one enriched in CD4 T cells and CD8 T cells, and another enriched in macrophages and neutrophils. Accordingly, we named them “lymphocyte-rich tumor” (cyan in **Fig. 5B**) and “myeloid cell-rich tumor” (purple in **Fig. 5B**). These two primary microenvironment archetypes were observed in relevance to exposed and interfaced LAs respectively (**Fig. 4C**), and they exhibited opposing indications for response or resistance. Consistent with findings from prior studies[17,26,49,50], lymphocyte-rich tumor microenvironments, which we observed near exposed LAs, were significantly more abundant among responders in pretreatment conditions (**Fig. 5C**). Conversely, myeloid cell-rich tumor microenvironments were predominantly present in nonresponder samples. Another tumor-associated microenvironment archetype characterized increased proportions of endothelial cells (“endothelial cell-rich tumor”, green in **Fig. 5B**), was observed primarily in nonresponders (**Fig. 5C**).

To characterize their distinctions in cellular organizations, we examined the abundance of non-tumor cells and cell-cell interactions (**Methods**). Lymphocyte-rich tumors exhibited higher densities of CD4 and CD8 T cells, with CD8 T cells engaging more directly with tumor cells (**Fig. 5D**, **Supplementary Fig. 16A,C**), consistent with proximity to exposed LAs. Dendritic cell-T cell interactions (**Supplementary Fig. 16B**) and immune cell “triads” (dendritic cells, CD4, and CD8 T cells), previously reported to increase immunotherapy efficacy[51,52], were also enriched (**Fig. 5E**). These trends were present in pretreatment samples, and became more pronounced following treatment (**Fig. 5E**, **Supplementary Fig. 16A,C**).

In contrast, myeloid cell-rich tumor microenvironments were characterized by CD68+/CD14+ macrophage presence. Replacing macrophages with T cells through *in silico* permutation and model evaluation (**Methods**) reversed the resistance indication, suggesting that our MIF-GNN model captured lymphocyte infiltration as a primary signature of response (**Supplementary Fig. 18**). In both resistance-associated microenvironment archetypes: myeloid cell-rich and endothelial cell-rich tumor microenvironments, enhanced tumor-endothelium interactions (**Fig. 5D**,**F**) and diminished immune triad presence (**Fig. 5E**) were observed.

We next compared and investigated gene expression programs of these microenvironment archetypes to reveal their underlying functional states (**Methods**). Unsurprisingly, spots mapped to lymphocyte-rich tumors exhibited significantly greater expression of known T cell activation genes (**Fig. 5G** top left, **Supplementary Fig. 14D**). Corresponding posttreatment neighborhoods showed a roughly four-fold additional increase. Elevated signals associated with interferon responses, inflammatory chemokines, and antigen presentation were also notable (**Supplementary Fig. 16D-E**). Importantly, we noticed enrichment of ECM disassembly signals in lymphocyte-rich tumors (**Fig. 5G** bottom right, **Supplementary Fig. 14G**), contrasting the ECM stabilization signatures seen in myeloid-rich tumor and in interfaced LAs (**Fig. 4G-H**). Together, these findings point to ECM disassembly and remodeling potential enablers of lymphocyte infiltration and activation during immunotherapy, while fibrotic ECM or inhibition of ECM disassembly may prevent successful therapeutic response.

Resistance-associated, myeloid-rich tumor microenvironments were further characterized by heightened expression of metabolism-associated genes (e.g., fatty acid, cholesterol, bile acid metabolism) (**Fig. 5G** top right, **Supplementary Fig. 14E**), as well as angiogenesis factors including transforming growth factor beta (TGFβ) and vascular endothelial growth factor (VEGF) signaling (**Fig. 5G** bottom left, **Supplementary Fig. 15D**, **Supplementary Fig. 16D**). These signatures varied across the patient cohort.

To determine the dynamics of gene expression during treatment, we compared corresponding ST spots from pretreatment and posttreatment samples (**Fig. 5H**, **Methods**) on the two more prevalent microenvironment archetypes. In lymphocyte-rich tumor microenvironments, genes highly expressed in pretreatment samples were associated with tissue identity, with transcription factors from various cell lineages (e.g., *GATA5*, *SOX18*, *LHX2*, *SOX2*), particularly neuronal lineages, upregulated. Following treatment, a dramatic increase in interferon-mediated immune response genes was observed, with *CXCL9* and *CXCL10* emerging as key chemokines (**Supplementary Fig. 16E**), and *CD38* as a key cell receptor. In myeloid-rich regions, fatty acid related metabolic genes (e.g., *CPS1, PCSK9*) were upregulated in pretreatment samples. Notably, some interferon-mediated immune genes (e.g., *IFI27*, *IRF1, IRF7*) exhibited high expression pretreatment, potentially indicating a preexisting inflammatory state. Posttreatment, genes associated with fibrotic ECM (e.g., *THBS1*, *CCN2, DCN, COL1A1*) formation became upregulated (**Fig. 5H**, **Supplementary Fig. 14F**). This observation, in resonance with the ECM signatures observed in interfaced LAs (**Fig. 4G-H**), suggest the development of a myeloid-associated ECM barrier in nonresponders, as opposed to the permissive ECM remodeling characteristic in response-indicative microenvironments discussed above.

Collectively, these findings highlight the distinct initial conditions—pre-existing T cell infiltration, angiogenic activities, dysregulated developmental states versus liver-like metabolic states—and the treatment-induced dynamics—immune activation driven by chemokines and cytokines, ECM-mediated immune exclusion—in tumor-associated microenvironments.

### Fingerprint of response and resistance across patients

Finally, we consolidated all the findings from the microenvironment-based analyses to curate a panel of spatially-resolved multi-omic scores. (**Methods**). This “fingerprint,” which we call SPARC (scoring pathways and architectures), incorporated cellular and transcriptomic signatures prioritized by our deep learning models and microenvironment-guided differential expression analyses. They include scores for LA structure (exposed versus interfaced), fibrotic or remodeling ECM gene expression states in LAs, T cell infiltration and activation within tumor microenvironments, metabolism programs, tumor-endothelium cell-cell interactions, and angiogenic pathway activity. Named gene expression signatures were curated based on differential expression analyses.

Each patient was assessed using the SPARC panel, and the averages for responders and non-responders were calculated. The results are presented as radial plots, with each axis representing the relative enrichment of a specific signature (**Fig. 6**, **Supplementary Fig. 19**). Immunotherapy responders exhibited strong immune activation profiles, characterized by greater intratumoral lymphocyte infiltration and higher levels of LA exposure. Transcriptomic signatures suggested treatment-induced intratumoral T cell activation, accompanied by heightened inflammatory chemokine expression, interferon responses, and ECM remodeling. Responder patients also exhibited a substantial decrease in metabolic expression signatures, potentially driven by necrosis of tumor and tumor-adjacent liver cells.

Conversely, nonresponders lacked sufficient intratumoral lymphocyte activities, and they displayed pronounced metabolism, angiogenesis, TGFβ, and immune-excluding fibrotic ECM signatures. Treatment-induced transitions—such as chemokine alterations, LA exposure increase, IFN-γ activation—were observed, though they were weaker and inconsistent across patients. Despite these overall trends, considerable heterogeneity on resistance signatures were observed across nonresponders (**Supplementary Fig. 19**), highlighting the need for personalized assessment of tumor microenvironment and treatment design for immunotherapy-resistant patients.

To connect our immunotherapy signatures to a broader oncology context, we asked whether any SPARC gene sets might be associated with differences in HCC outcomes more generally. We obtained per-gene expression hazard ratios derived from The Cancer Genome Atlas (TCGA) data[80]. In this dataset, higher scores mean that higher expression of a gene is associated with increased risk of death and poor prognosis. We also calculated spot-level correlations between gene expression in our ST data and the ST-MLP model predictions to derive a metric for associating individual genes with immunotherapy response. Surprisingly, many genes that our model associated with nonresponse to immunotherapy predicted better prognoses outside the immunotherapy context (**Supplementary Fig. 20A**). Moreover, several SPARC nonresponse signatures had a strong bias towards lower hazard ratios and better prognoses, while response signatures were more neutral (**Supplementary Fig. 20B**). We hypothesize that certain tissue factors that slow cancer metastasis or invasion may also inhibit immune infiltration and activation.

## Discussion

In this study, we analyzed spatially-resolved, multi-omic profiling of hepatocellular carcinoma samples collected before and after perioperative immunotherapy. Through multimodal integration followed by interpretable deep learning modeling, we identified key microenvironment archetypes. These archetypes revealed the cellular and transcriptomic characteristics of immune and tumor-intrinsic niches, as well as their dynamic changes during immunotherapy. By consolidating these findings, we developed SPARC, a comprehensive fingerprint to capture the landscape of HCC tumor microenvironment undergoing immunotherapy. This fingerprint offers holistic insights into the underlying mechanisms of immunotherapy outcomes and helps pave the way to precision oncology in HCC.

To achieve our analysis goal, we developed a computational strategy to integrate the complementary powers of MIF and ST: we used MIF-GNN to characterize detailed spatial-cellular microenvironments and to organize them into archetypes that have quantifiable effects on treatment response. We then used ST to uncover expression and pathway signatures linked to each response- or resistance-associated archetype. Importantly, our strategy is extensible and generalizable to other disease-centric studies, where neighborhood-scale determinants of key disease characteristics (e.g., prognosis, treatment response) can be extracted and analyzed. We anticipate that, as more spatially-resolved, multi-omic datasets are generated, our approach will facilitate discovery of new disease-relevant determinants and enable in-depth understanding of their signatures and underlying mechanisms.

In agreement with current literature, we show that pre-existing intratumoral T cell immunity is a critical precursor for response to immunotherapy in HCC. Lymphocyte-rich tumor microenvironments in pretreatment samples, characterized by high T cell infiltration and immune triad formation, were strongly associated with treatment response. Similar correlations have been reported in prior studies on HCC[17], head-and-neck cancer[53], and melanoma[20]. Furthermore, we dissect comparisons of lymphocyte-rich tumor microenvironments between pre- and posttreatment samples to reveal the specific T cell activation programs, elevated interferon responses[18], inflammatory chemokines (*CXCL9/10*), and ECM disassembly and remodeling programs that dynamically respond to treatment.

We also create a novel classification for LAs of distinct structures in tumor-adjacent stroma. Previous studies observed that LA localization[54] and TLS organization[44] may impact immunotherapy response. In our analysis, exposed LAs, which come into direct contact with adjacent tumors, were identified as strong response indicators. They exhibited greater B cell and CD8 T cell presence, higher expression of Th1- and Tfh-associated[55] proteins and genes, and elevated interferon gamma signaling and *CXCL13* expression. These features could conceivably represent the early formation of functional TLS, known contributors to immunotherapy response[43,45,56,57]. We also observed a contrasting group of interfaced LAs that was almost exclusive to nonresponders. These LAs are defined by a unique interface niche that physically separates the LA from neighboring tissue. By differential expression analysis, we found that a restrictive ECM characterizes the barrier between these nascent immune structures and tissue[58–60], which was also reflected in their nearby tumor-associated microenvironments. Computational permutations, supported by some previous studies on key relevant genes[21,61] suggest that interventions to degrade these interfaces could sensitize a resistant tumor microenvironment to immunotherapy.

The most consistent signatures of nonresponse within the tumor microenvironment itself were myeloid cell infiltration and elevated liver-associated metabolism programs, both known correlates of tumor progression and drug resistance[62,63]. Adding to these reports, we curated signatures of chemokines and immune-excluding fibrotic ECM genes that are expressed in immunotherapy-resistant patient tumors. We also identified angiogenesis, tumor-endothelium interactions, and TGFβ signaling as negative correlates of immunotherapy response, which are all well-documented contributors to immunosuppressive tumor microenvironments[31,64,65]. Strikingly, in most patients, these conditions do not meaningfully change during checkpoint inhibition. Joint therapies (e.g. anti-VEGF plus immune checkpoint inhibitor[66], anti-TGFBR1 and anti-PDL1[31]) and therapies targeting myeloid-derived immune suppression[67] and fatty acid metabolism[68–70] have been explored in the context of immunotherapy, and the holistic cell and expression “fingerprints” we have defined here may aid in personalizing such treatment combinations.

Collectively, this study describes the complex and highly heterogeneous landscape of HCC, highlighting key signatures for both response and resistance to immunotherapy. Our findings suggest a number of cellular and molecular targets for further therapeutic development, and, together with other recent works[17,21,31,70], point to the promising approach of addressing immunotherapy resistance by jointly applying antagonists targeting metabolism and other pathways. Further efforts to stratify patients based on their tumor-intrinsic and immunological profiles could enable personalized design of such combination therapies, improving clinical outcomes.

Last, we note several limitations to the current study: First, our cohort was limited to 13 patients, so further validation of the signatures on larger cohorts will be important for generalization. Second, although our analysis handles the differences in resolution between MIF and ST data, the current study was limited by spot-level resolution of ST. Further advances in spatial profiling techniques or computational tools could increase the accuracy of our results and interpreations[71]. Third, although our cohort included patients treated with anti-PD1 and combination anti-PD1 and anti-CTLA4, our study lacks the numbers to meaningfully distinguish these subgroups. Further work will extend our findings to additional patient populations and oncology contexts. Altogether, this work underscores the value of spatially-resolved multi-omics for immuno-oncology and opens up opportunities to improve precision immunotherapy through integrative, spatially-resolved patient profiling.

## Data and code availability

The processed spatial multi-omics data and code used in this study are publicly available through https://doi.org/10.5281/zenodo.15392699. This repository includes:

● **Processed CODEX MIF Data**: Centroid locations, normalized protein biomarker expression levels, and cell type annotations for all segmented cells across 48 fields of view.
● **Processed Visium ST Data**: Gene expression profiles and co-registered spatial coordinates for all spots across 22 samples.
● **Data Analysis Notebook**: A fully documented Jupyter Notebook containing the workflows and code used to generate all figure panels in the manuscript.

## Methods

### Spatial transcriptomics assay (10X Genomics, Visium)

Formalin-fixed paraffin-embedded (FFPE) tissue from HCC samples were used for spatial transcriptomics analysis. HCC tumors were paraffin embedded and serially sectioned (thickness 5 μm). FFPE samples were tested for RNA quality with an DV200 > 30% (Agilent). The samples were then processed according to the standard Visium Spatial Gene Expression protocol (10x Genomics) using the Visium Spatial Gene Expression Slide & Reagent Kit (10x Genomics). Libraries were cleaned up using SPRI select reagent and quantified using the High Sensitivity DNA Kit run on Agilent 2100 Bioanalyzer and also KAPA Illumina library quantification kit (Roche, 7960140001) run on LightCycler 480. The library pool was quantified on Bioanalyzer and with quantitative PCR and sequenced using Illumina NextSeq 500.

### Spatial proteomics/multiplexed immunofluorescence assay (CODEX)

4 µm FFPE tissue sections from all samples were placed on a positively charged glass slide and stored at 4°C until use. CODEX staining assay was performed following the manufacturer’s protocol. Purified antibodies, barcodes and reporters were purchased commercially as listed in **Supplementary Table 2**. Antibodies CD1c, CD209, CXCR5, PD-1, CD56 and Podoplanin were conjugated through the Spatial Tissue Exploration Program (STEP) from Akoya Biosciences and Leinco Technologies. For another 23 biomarkers, purified antibodies (free of BSA and glycerol) were conjugated in-house. The remaining barcode conjugated antibodies were purchased commercially from Akoya Biosciences. All purified barcoded-conjugated antibodies were used within 6 months of conjugation. The stained tissue section was captured using PhenoCycler-open with Keyence microscope using DAPI, ATTO550, CY5 and AF488 filters at a scan resolution of 0.50 µm (20X). Alignment of images across cycles, stitching of tiles and subtraction of auto-fluorescence was performed using CODEX® Processor application. Cells were segmented by applying StarDist[72], a deep learning-based algorithm, to the DAPI channel.

### Quality control and normalization of ST data

We excluded spots for which there were less than 100 counts, and genes that were present in less than 10 spots (**Supplementary Fig. 1D-E**). After quality control, we used the package `scanpy` to perform basic downstream analysis. In particular, gene expression counts were normalized per spot by scaling to counts per million, and then scaled using a log1p transform. For differential analysis, we retained the top 5000 highly expressed genes using the Seurat v3 method[73].

### Cell type clustering of MIF data

To annotate cell types, we obtained a cell-by-protein biomarker expression matrix by integrating over all the channels of the MIF data. We then constructed a k-nearest neighbour graph (k = 75) on the expression matrix and performed Leiden graph clustering[74]. Clusters were manually annotated according to their protein biomarker expression patterns. Average expression on the full biomarker panel of each cell type is visualized in **Supplementary Fig. 4A**. UMAP dimensionality reduction[75] of the biomarker expression matrix colored by cell type annotation is visualized in **Supplementary Fig. 4B**. Outlying clusters that contain cells located in imaging artifacts were manually examined and annotated as “Unknown/Uncharacterized” (**Supplementary Fig. 1A-C**). These cells were excluded from most of the downstream analysis.

### Alignment and pseudo-spotting

We aligned ST samples with their corresponding MIF samples (DAPI stains) using the H&E images collected during the experiment (**Supplementary Fig. 1E**). The alignment process employed the Scale-Invariant Feature Transform (SIFT) algorithm[76,77], which identified key points in images and matched the feature descriptors of key points to identify corresponding pairs of points from different images. Based on the matched points between MIF and ST-derived H&E, a transformation was performed to map the coordinate system of the ST samples to that of the MIF samples.

We first converted the H&E images into grayscale and adjusted their contrast using Contrast Limited Adaptive Histogram Equalization (CLAHE; implemented in OpenCV). The SIFT algorithm was then applied to detect features in both the H&E and DAPI images. Feature matching was performed based on the SIFT descriptors, with manual adjustments to the feature matching ratio depending on the similarity of the images. A transformation matrix was estimated using the Random Sample Consensus (RANSAC) algorithm, allowing for rotation, translation, and scaling transformations. The RANSAC inlier threshold was also manually adjusted to accommodate varying degrees of distortion between images.

With the matched coordinate system, we identified spatially-overlapping sets of cells for ST spots using a distance-thresholded K-nearest neighbor approach. In particular, for each spot, its K nearest cells (K = 50) identified in the MIF data are extracted and filtered by a maximum distance (R = 100 µm). This process is referred to as “pseudo-spotting” in the text. Subsequently, using the annotations assigned to MIF cells (e.g., cell type or microenvironment archetype), we further identified ST spots highly “enriched” in a specific annotation class, defined as >75% of neighboring cells sharing the annotation. These enriched spots were collected as representatives of the target annotation class.

### Copy number alteration analysis and malignancy scores

Copy number alterations were assessed in patient samples using InferCNV[39]: we employed a normal human liver cell atlas[78] as the reference for normal cells, and treated ST spots as individual units, from which the raw gene counts were assembled to construct the input matrix.

InferCNV was then run using default parameters, with cells from the normal liver atlas serving as the reference group (i.e., annotated as “normal”) and ST spots as query (i.e., annotated as “malignany_{compartment}”). Following the inference of CNA matrices, we calculated a malignancy score for each ST spot by averaging the deviations, including both amplification and deletion, across all genes, serving as a measure of genomic instability and mutation burden.

### Predictive Modeling

To identify key features associated with immunotherapy response or resistance, we conducted a predictive modeling study using MIF and ST data to predict patient-level responses to treatment. This experiment was conducted using a four-fold cross-validation approach: patients were divided into four groups, each group containing 1–2 responders and 1–2 nonresponders. In each validation fold, one group was reserved for evaluation while the remaining groups were used for training. Predictions for the test patients were collected and later aggregated across all four folds to compute accuracy metrics. This validation scheme was applied consistently across all methods described below.

#### Sample-scale baselines

We used MIF- and ST-based sample-scale methods as baseline models. For the MIF data, each sample was featurized as a composition vector of cell types. For the ST data, pseudo-bulk RNA-seq samples were generated by randomly combining 200 ST spots, mimicking bulk RNA sequencing. These features were then used as inputs to standard machine learning models, including logistic regression and XGBoost classifiers, to predict patient-level treatment responses.

#### MIF-GNN

For MIF data, we employed a previously developed graph neural network (GNN) approach[30] to encode neighborhoods of cells. These neighborhoods were defined as 3-hop subgraphs extracted from the Delaunay triangulated graph representation of the MIF region/FOV. The GNN uses discrete cell types and neighborhood graphs as inputs, and employs a 3-layer graph isomorphism network to generate embeddings for neighborhoods, which are further passed through a multi-layer perceptron (MLP) predictor to infer patient-level labels: response/resistance to immunotherapy.

During inference, each test region was first divided into collections of neighborhoods (i.e., 3-hop subgraphs). The trained GNN was then used to independently generate predictions for each neighborhood. The resulting set of prediction values were averaged to yield the aggregated prediction for the test region.

#### ST-MLP

Informed by the comparable spatial dimensions of MIF neighborhoods (50-100 µm diameter) and ST spots (55 µm), we conducted a paralleled predictive modeling analysis on ST data. We applied Harmony[79] on ST spots to extract 200 batch corrected principal components for use as features. We then trained MLP models on these features to predict patient-level labels for 500 epochs using the Adam optimizer with a learning rate of 1e-3 and early stopping.

During inference, predictions were similarly collected across all spots from the test sample and averaged to yield the aggregated prediction. To compare predictions generated from ST and MIF data at the scale of neighborhoods, we performed pseudo-spot aggregation over MIF-based predictions from corresponding cells and compared the aggregated values with ST-based predictions.

### Identification of microenvironments

Clusters of neighborhoods, or microenvironment archetypes, were identified through clustering of neighborhood embeddings[30] generated by a GNN model trained on predicting immunotherapy response using all pretreatment MIF samples:

– We first randomly sampled a reference set comprising a total of 50,000 neighborhoods and calculated their neighborhood embeddings using the trained GNN model;
– We then applied Leiden clustering on the reference embeddings to derive clusters of neighborhoods. The resolution parameter was tuned to balance granularity, polarity (if clusters are strongly associated with response or resistance), and universality (if the clusters are present in more than one sample/patient);
– We also calculated UMAP dimensionality reduction of the top 20 principal components of reference embeddings and generated 2D visualizations of the clusters and embedding space (**Supplementary Fig. 7A**);
– We trained an XGBClassifier using the reference embeddings and their neighborhood clusters to derive a multi-class embedding-to-microenvironment predictor;
– We collected embeddings for all neighborhoods across all pretreatment and posttreatment samples and applied the embedding-to-microenvironment predictor to collect microenvironment archetype annotations for all cells/neighborhoods.

### Analysis of cellular organization via *in silico* permutations

To investigate differences in cellular organizations across different microenvironment archetypes, we performed *in silico* permutations on subgraphs and evaluated the influences of such permutations on predicted probabilities of responding and microenvironment archetype annotations. Two major types of permutations were performed:

– We altered the identities of the target cell(s) within a subgraph of interest without modifying other cells or the underlying graph structure. For instance, in pretreatment myeloid cell-rich tumor microenvironments, we randomly replaced myeloid cells with lymphocytes (**Supplementary Fig. 18**);
– We extracted all cells from a subgraph of interest and re-ordered them to simulate different levels of mixing between target cell type(s). In particular, we first transformed the coordinates of all cells from a subgraph into polar coordinates with regard to the subgraph center, then sorted them based on their azimuth angles. We next sorted the cell identities (i.e., cell types) in a specific order so that pre-specified groups of cells are mixed together while being separated from other cell types. The re-ordered list of cell identities were reassigned to the nodes in the subgraph in the order of azimuth angles. The resulting permuted subgraph has identical cell type composition but altered spatial organizations (**Supplementary** Fig. 12A-C).

After permutations, we used the same GNN model and embedding-to-microenvironment predictor to infer the permuted microenvironment archetype annotations and predictions for immunotherapy response.

### Differential expression and gene set enrichment analysis

We applied pseudo-spotting on microenvironment archetype annotations and identified representative spots that are highly enriched in one type of microenvironment. To characterize their associated gene signatures, we performed differential expression analysis on the normalized gene expression values of representative spots using t-test. Expression of these spots are compared against a reference set of spots to identify differentially expressed genes (DEGs). After ranking these genes using the test statistic, we optionally ran GSEA on the Gene Ontology Biological Process (GOBP) and Molecular Signatures Database Hallmark (MSigDb Hallmark) gene sets to extract key molecular programs.

The one-versus all DEG analysis was conducted by comparing microenvironment-specific spots against all other spots (**Supplementary Fig. 9**). Comparative analysis between two microenvironment archetypes used one group as the query and the contrasting group as reference to identify DEGs.

### Characterization of microenvironment archetypes

To characterize the distinctions between microenvironment archetypes, or dynamics in the same microenvironment archetype after treatment, we performed targeted comparisons between groups of neighborhoods and their corresponding ST spots. Various cellular and transcriptomic features were assessed.

#### Cellular features

We assessed differences in spatial cellular organization across neighborhood groups along the following axes:

– Compositions of cell types (e.g., immune cells) in the neighborhoods;
– Compositions of the neighboring cells of a specific cell type of interest (e.g., tumor cells) in the neighborhoods. Neighboring pairs of cells were defined based on the graph representation of MIF region generated via Delaunay triangulation;
– Enrichment of interactions between cell types, measured by the observed counts of neighboring cell pairs of specific types divided by the expected counts if cells are randomly placed in the neighborhood. We further showed the differences in enrichment matrix to highlight alterations in cell-cell interaction patterns.

#### Transcriptomic features

We also explored differences in transcriptomic profiles using corresponding ST spots via:

– Differential expression analysis between two contrasting groups of ST spots (e.g., lymphocyte-rich versus myeloid cell-rich tumor spots, lymphocyte-rich tumor spots from pretreatment samples versus posttreatment samples);
– Targeted evaluations on key gene markers or programs informed by cellular features or DEG results. For example, we used the GOBP term: *T Cell Activation Involved In Immune Response (GO:0002286)* to assess intratumoral T cell activations in lymphocyte-rich and myeloid cell-rich tumor microenvironments. In the LA analysis, we used known gene markers to assess the activities and/or abundance of different T cell subtypes.

#### Annotation and categorization of LAs

Using human annotations of LAs as reference, we identified four microenvironment archetypes uniquely associated with LAs (**Supplementary Fig. 7B, Fig. 3E, Fig. 4A**)): CD4 T rich LA, B rich LA, tumor interface at LA rim, and fibroblast interface at LA rim.

To systematically identify all LAs in the data, we first combined the two core microenvironment archetypes (i.e., CD4 T rich LA and B rich LA) and extracted all spatially connected components containing more than 15 cells. These connected units were designated as LA cores. We then expanded each LA core to include surrounding tumor and fibroblast interface neighborhoods. This was done iteratively by examining neighboring cells and adding those whose neighborhoods were annotated as tumor interface or fibroblast interface. The expansion was repeated for up to six iterations or until no additional neighboring cells were added. Examples of the resulting annotations are shown in **Fig. 4D**.

Once all LAs were identified, we classified them as either “exposed” or “interfaced” based on the presence and proportion of interface microenvironments. Specifically, LAs with >50% of neighborhoods in the core categories (CD4 T rich LA or B rich LA) were labeled as *exposed*. Among the remaining LAs, those in which tumor interface microenvironments comprised more than one-third of the non-core neighborhoods were labeled as *interfaced*.

### Curation of response and resistance signatures

To assess response and resistance-associated signatures at patient level, we curated a panel of signatures based on microenvironment-based analyses. Each signature was calculated using all neighborhoods or spots from an individual patient.

#### MIF-based signatures

Following signatures were calculated based on the cell type and microenvironment archetype annotations of cells segmented in MIF regions.

– **Intratumoral lymphocyte presence** — defined as the ratio of lymphocyte-rich tumor microenvironments to the sum of lymphocyte-rich and myeloid cell-rich tumor microenvironments;
– **Tumor-endothelium interaction** — defined as the ratio of endothelial cells located in myeloid cell-rich and endothelial cell-rich tumor microenvironments—where heightened tumor-endothelium interaction is observed—to the total number of endothelial cells across all four tumor-associated microenvironments.
– **LA exposure level** — calculated as the ratio of core LA microenvironments (CD4 T rich LA and B rich LA) to the sum of core LA and tumor interface microenvironments.

#### ST-based signatures

To address the limited number of spots with direct pseudo-spot matches, we first trained two auxiliary machine learning models (XGBClassifier) to predict the compartment (e.g., tumor, stroma, tumor-adjacent liver) and the major microenvironment class of each ST spot based on the normalized expression of highly variable genes. These models were trained on the subset of spots with known compartment and microenvironment archetype annotations derived from pseudo-spotting, and subsequently applied to infer labels for all remaining spots.

The following signatures were then computed using the normalized expression of genes of interest in spots assigned to specific compartments or microenvironments via either pseudo-spotting or the auxiliary predictors:

– **Intratumoral T cell activation** — defined as the aggregated expression of genes associated with two GOBP terms: Antigen Processing And Presentation Of Peptide Antigen Via MHC Class II (GO:0002495) and T Cell Activation Involved In Immune Response (GO:0002286), calculated in spots mapped to tumor-associated microenvironments.
– **IFN-γ response** — defined as the aggregated expression of genes associated with two GOBP terms: *Cellular Response To Type II Interferon (GO:0071346)* and *Type II Interferon-Mediated Signaling Pathway (GO:0060333)*, calculated in spots mapped to LAs (CD4 T rich LA, B rich LA, tumor interface and fibroblast interface) and lymphocyte-rich tumor microenvironments.
– **Positive angiogenesis regulation** — defined as the aggregated expression of a set of 13 genes (**Supplementary Table 3**), identified as the leading-edge subset of the GOBP term *Positive Regulation of Angiogenesis (GO:0045766)* from the GSEA analysis between lymphocyte-rich and myeloid cell-rich tumor microenvironments, calculated in spots mapped to all tumor-associated microenvironments.
– **Metabolism strength** — defined as the aggregated expression of a set of 75 genes (**Supplementary Table 3**), identified as leading-edge subsets of five metabolism-related GOBP terms: *Fatty Acid Metabolic Process (GO:0006631)*, *Steroid Metabolic Process (GO:0008202)*, *Monocarboxylic Acid Metabolic Process (GO:0032787)*, *Bile Acid Metabolic Process (GO:0008206)*, and *Arginine Metabolic Process (GO:0006525)*, from the GSEA analysis between lymphocyte-rich and myeloid cell-rich tumor microenvironments, calculated in spots mapped to all tumor-associated microenvironments.
– **TGFβ signaling and response** — defined as the aggregated expression of a set of 25 genes (**Supplementary Table 3**), identified as leading-edge subsets of the GOBP terms *Transforming Growth Factor Beta Receptor Signaling Pathway (GO:0007179)* and *Cellular Response To Transforming Growth Factor Beta Stimulus (GO:0071560)* from the GSEA analysis between lymphocyte-rich and myeloid cell-rich tumor microenvironments, calculated in spots mapped to all tumor-associated microenvironments.
– **Inflammatory chemokines** — defined as the aggregated expression of two sets of chemokines (**Supplementary Table 3**). Both sets were identified through the targeted chemokine analysis in tumor-associated microenvironments (**Supplementary** Fig. 16E). This signature was calculated as the difference between inflammatory set and homeostatic set chemokine expression in spots mapped to all tumor-associated microenvironments.
– **Immune-excluding ECM and remodeling/disassembly** — defined as the aggregated expression of two sets of genes (**Supplementary Table 3**). The first set, comprising 30 genes,was identified as a leading-edge subset of the GOBP term *Extracellular Matrix Organization (GO:0030198)*, derived from GSEA analysis between pretreatment and posttreatment spots of myeloid cell-rich tumor microenvironments. This set primarily includes structural ECM components such as collagens, fibronectins, and laminins, along with regulators of ECM assembly and stabilization. The second set, comprising 29 genes, was identified as subsets of the GOBP terms *Extracellular Matrix Organization (GO:0030198)* and *Extracellular Matrix Disassembly (GO:0022617)*, selected from DEGs and GSEA analysis between pretreatment and posttreatment spots of lymphocyte-rich tumor microenvironments. This set is enriched for ECM-degrading enzymes, including matrix metalloproteinases (MMPs) and ADAMTS proteases, as well as ECM-modifying factors involved in remodeling. The two sets of genes were calculated in spots mapped to tumor-associated microenvironments.

### Declaration of generative AI and AI-assisted technologies in the writing process

During the preparation of this work, the authors used GPT-4o to proofread and polish certain passages in the text. After using this tool, the authors reviewed and edited the content as needed. They take full responsibility for the content of the publication.

## Acknowledgements

The HCC clinical trial was supported via A.O.K. by The MD Anderson Cancer Center SPORE in Hepatocellular Carcinoma Grant P50 CA217674 and research funding from BMS. J.Z. is supported by funding from the Chan-Zuckerberg Biohub. We thank the software engineers at Enable Medicine for their support and contributions to compute infrastructure and data interactivity throughout the project.

## Author Contributions

A.E.T., J.Z., Z.W., A.T.M., and P.S. conceptualized the research. A.O.K. led the HCC clinical trial and clinical follow-up. M.L. analyzed the clinical trial. Z.W. led the overall project and performed the primary computational analysis, MIF model development, and data interpretation. J.B. led the ST analysis and ST model development. S.H. and J.B. performed ST data processing. S.J. and S.B. led the data collection, annotated the data, and guided data interpretation. M.B. developed the data co-registration approach. P.S., A.T.M., J.Z., and A.E.T. supervised the project. Z.W., J.B., and A.E.T. wrote the manuscript with contributions from all authors, and all authors approved the final version.

