## Supplementary Figures and Tables for "Spatial multi-omics and deep learning reveal fingerprints of immunotherapy response and resistance in hepatocellular carcinoma"

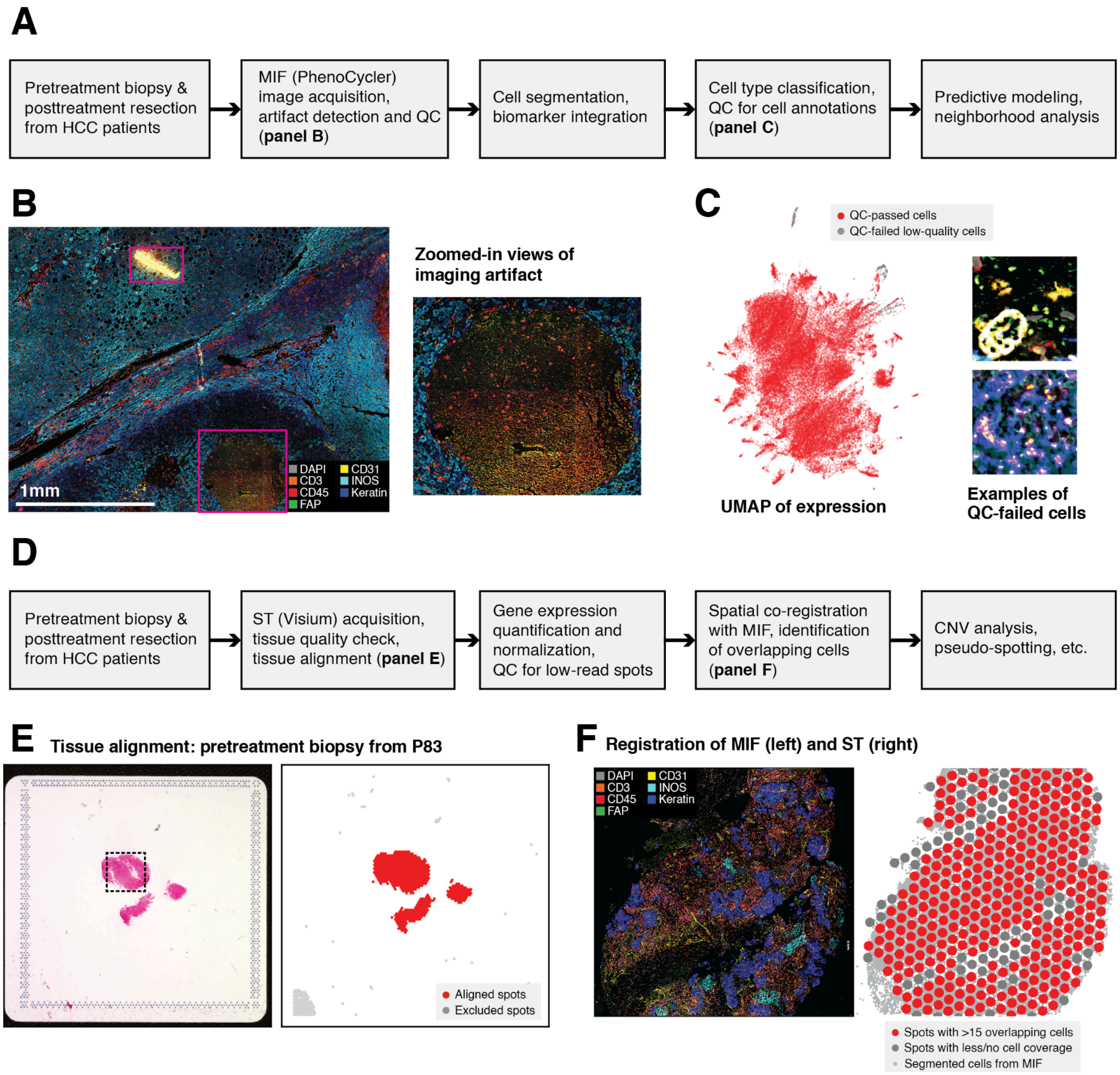


**Supplementary Fig. 1: Data pre-processing and quality control (QC) of MIF and ST data.**

**A.** MIF data processing pipeline. **B.** An MIF image with noticeable imaging artifacts, marked by magenta rectangles. Two FOVs are excluded from downstream analysis due to artifacts or failure during cell annotation. **C.** Cell populations with abnormal expression intensities (grey dots in the UMAP) detected during cell type classification. These cells are visually inspected and excluded through the cell annotation QC step. **D.** ST data processing pipeline. **E.** Example tissue alignment for a pretreatment biopsy sample. Spots not aligned to tissue (grey) are excluded from downstream analysis. **F.** Spatial registration of MIF and ST data from the same biopsy sample (dashed black square marked in panel **E**). ST spots with overlapping MIF cells (red) are used for pseudo-spotting and cross-modal analysis.

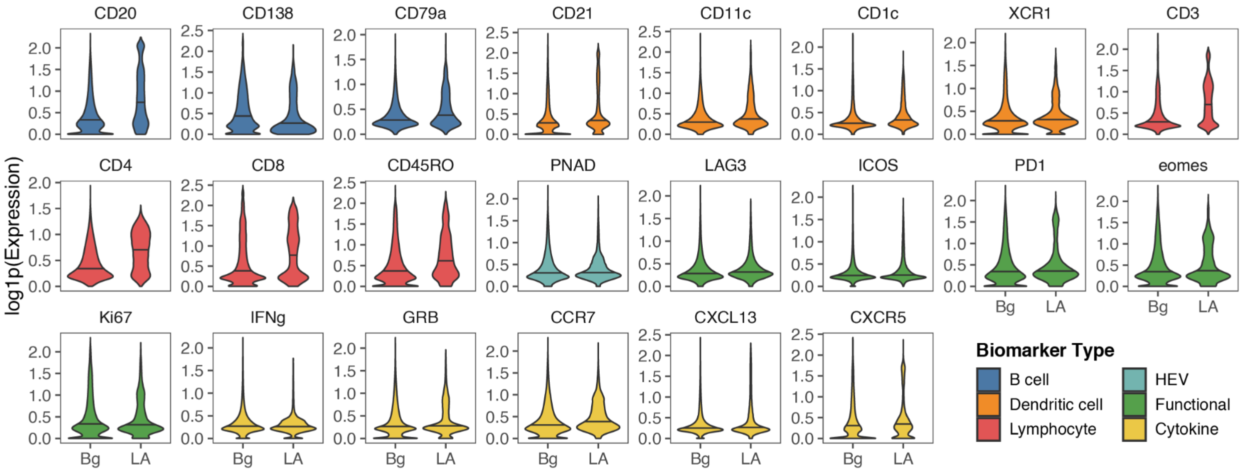


**Supplementary Fig. 2: Protein expression characteristics of LAs in HCC samples.**

Expression of proteins within pathologist-annotated LAs. Colors indicate the canonical category each marker is most associated with. HEV: high endothelial venule. Bg: background–non LA regions.

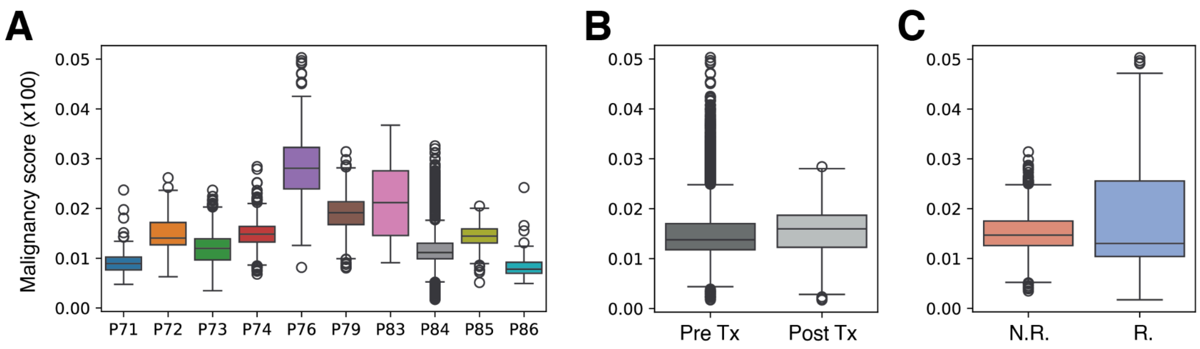


**Supplementary Fig. 3: Malignancy scores across groups.**

Malignancy scores of spots spatially co-registered to tumor and tumor-adjacent liver compartments are visualized according to **A.** patient, **B.** treatment state, and **C.** response to immunotherapy.

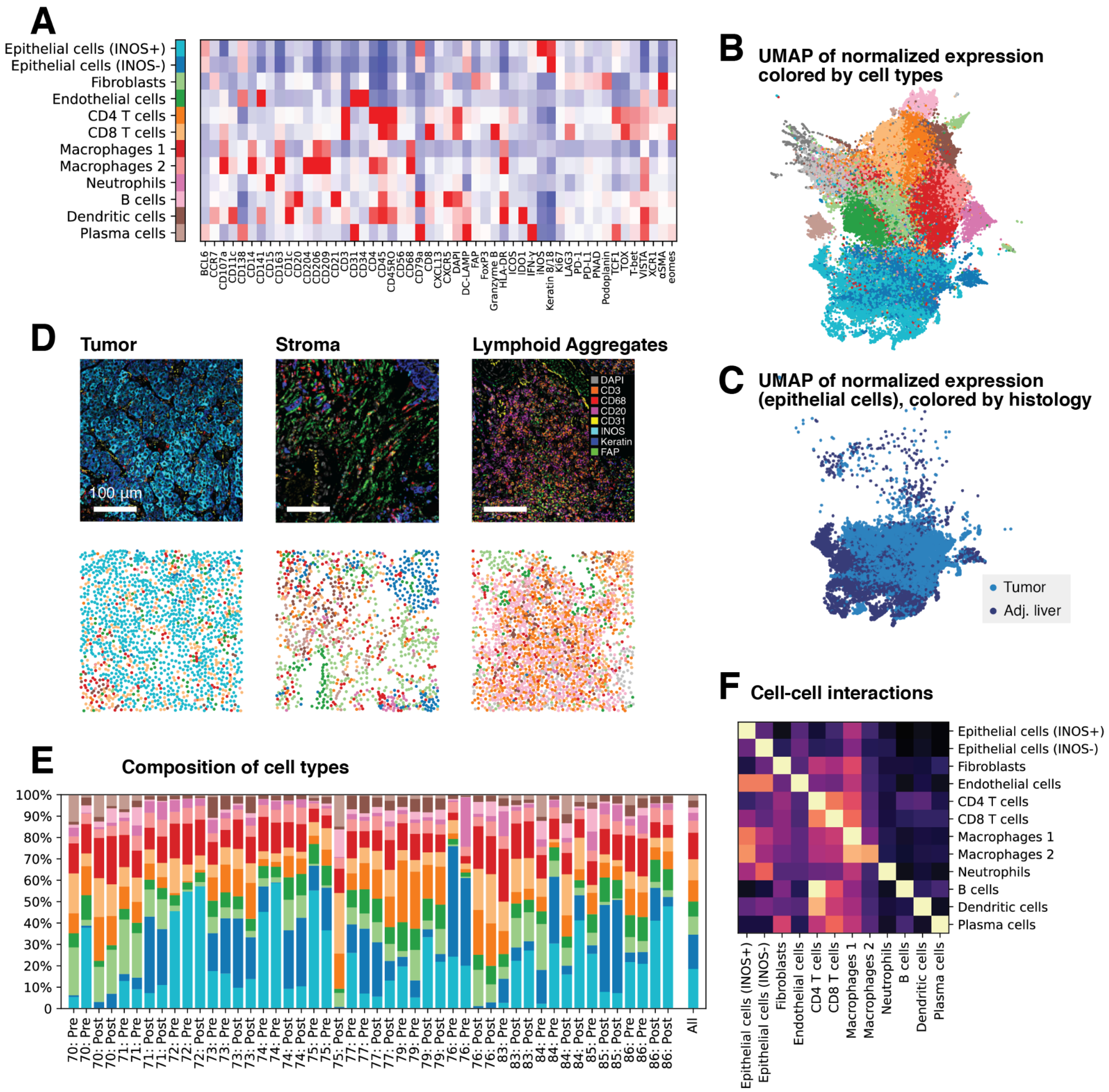


**Supplementary Fig. 4: Protein expression in cells define distinct cell types**

**A.** Heatmap showing the full expression profiles of measured protein biomarker expression across different cell types. **B.** Dimensionality reduction of cellular protein expression using UMAP. Colors represent different cell types (same as panel **A**). **C.** Epithelial cells from tumor and tumor-adjacent liver compartments marked in UMAP. They have similar protein expression profiles. **D.** Zoomed-in views of tumor and stroma compartments. Distinct spatial compositions and distributions of cells are observed. **E.** Compositions of cell types across samples. Note that cell type compositions within regions are not good predictors of responses to immunotherapy. **F.** Cell-cell interaction strengths. The heatmap highlights the major tumor-infiltrating immune cells and the co-localization patterns of different immune cells.

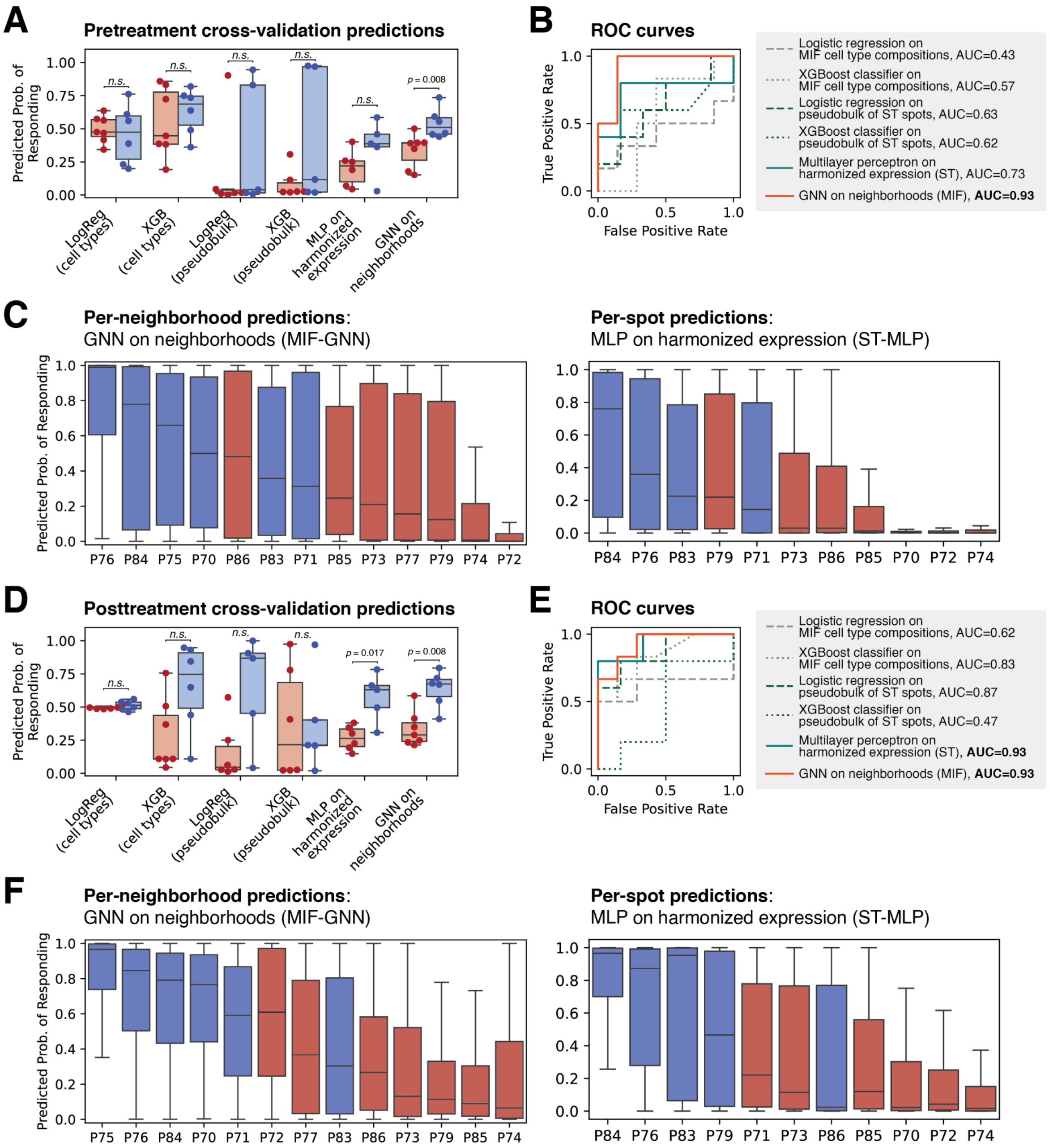


**Supplementary Fig. 5: Predictive modeling of responses to immunotherapy by cross-validation on pretreatment and posttreatment samples**

**A.** Test-time predictions from four-fold cross-validation on pretreatment samples. The box plots show mean predictions for each patient categorized by MPR. Across all benchmarked methods, neighborhood-scale methods achieve superior performances, in which MIF-GNN significantly differentiates nonresponders from responders (MWU test, $p = 0.008$). **B.** Receiver operating characteristic (ROC) curves of model predictions on pretreatment samples. Neighborhood-scale predictors (solid curves) outperform sample-scale predictors (dashed and dotted curves). MIF-GNN achieved the best AUROC of 0.93. **C.** Distributions of all neighborhood/spot-scale predictions on pretreatment samples. **D.**

Test-time predictions from four-fold cross-validation on posttreatment samples. MIF-GNN and ST-MLP differentiate responders from nonresponders with statistical significance (MWU test, $p = 0.017$ and 0.008). **E.** ROC curves of posttreatment predictions. Better performances are observed, in which MIF-GNN and ST-MLP exhibit the best performances. **F.** Distributions of all neighborhood/spot-scale predictions on posttreatment samples.

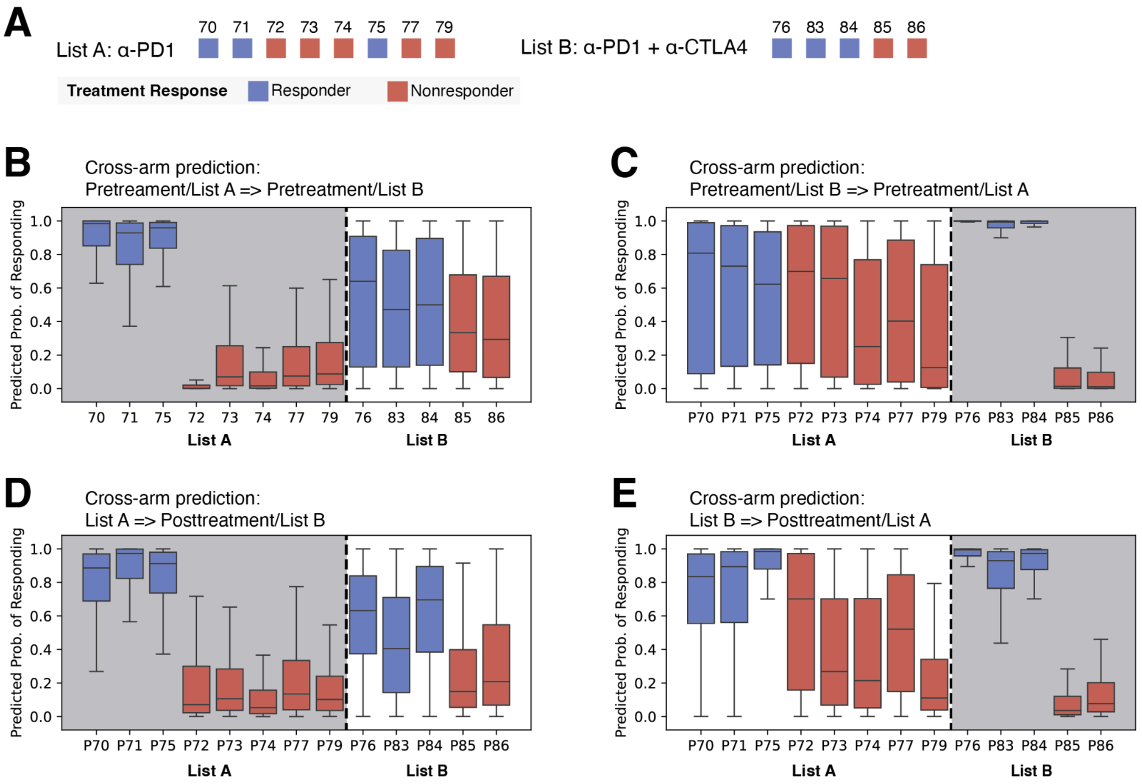


**Supplementary Fig. 6: Cross-treatment prediction suggests potential benefit of anti-CTLA4 treatment.**

**A.** The patient cohort received two types of treatment: list A patients received only anti-PD1 (nivolumab) treatment, while list B patients received a combination of anti-PD1 and anti-CTLA4 (ipilimumab) treatments. In the main text, we combined the two groups for larger-scale analyses. **B.** MIF-GNN model trained on list A patients' data is applied to list B patients. The box plots show the distributions of all neighborhood-scale predictions. Note that boxes within the shaded area are training samples, where better separation is expected. List B responders exhibited higher predicted probabilities of response than nonresponders. **C.** In a complementary experiment, MIF-GNN model trained on list B patients' data is applied to list A patients. Notably, patients P72 and P73, who were nonresponders to monotherapy (anti-PD1), exhibit high predicted probabilities of response to the combination treatment, suggesting a potential benefit of including anti-CTLA4 treatment for these patients. **D-E.** MIF-GNN models trained on both pretreatment and posttreatment data are evaluated for cross-treatment predictions. Prediction results largely agree with the observed response/resistance labels.

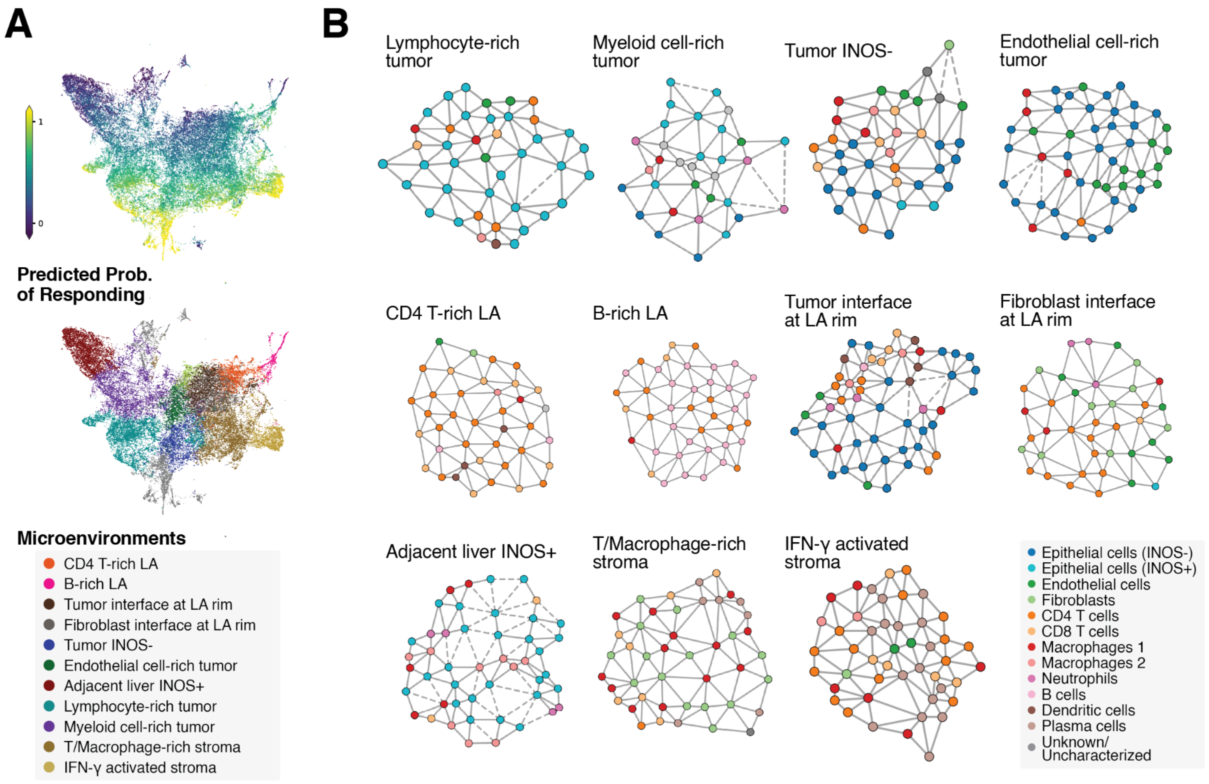


**Supplementary Fig. 7: Examples of microenvironment archetypes**

**A.** Dimensionality reduction of GNN embeddings via UMAP. A gradient of predicted probabilities of responding is revealed. An additional UMAP plot is shown with microenvironment archetype annotations. **B.** Example neighborhoods from major microenvironment archetypes. In these graphical representations, nodes represent cells, and edges represent neighboring pairs of cells.

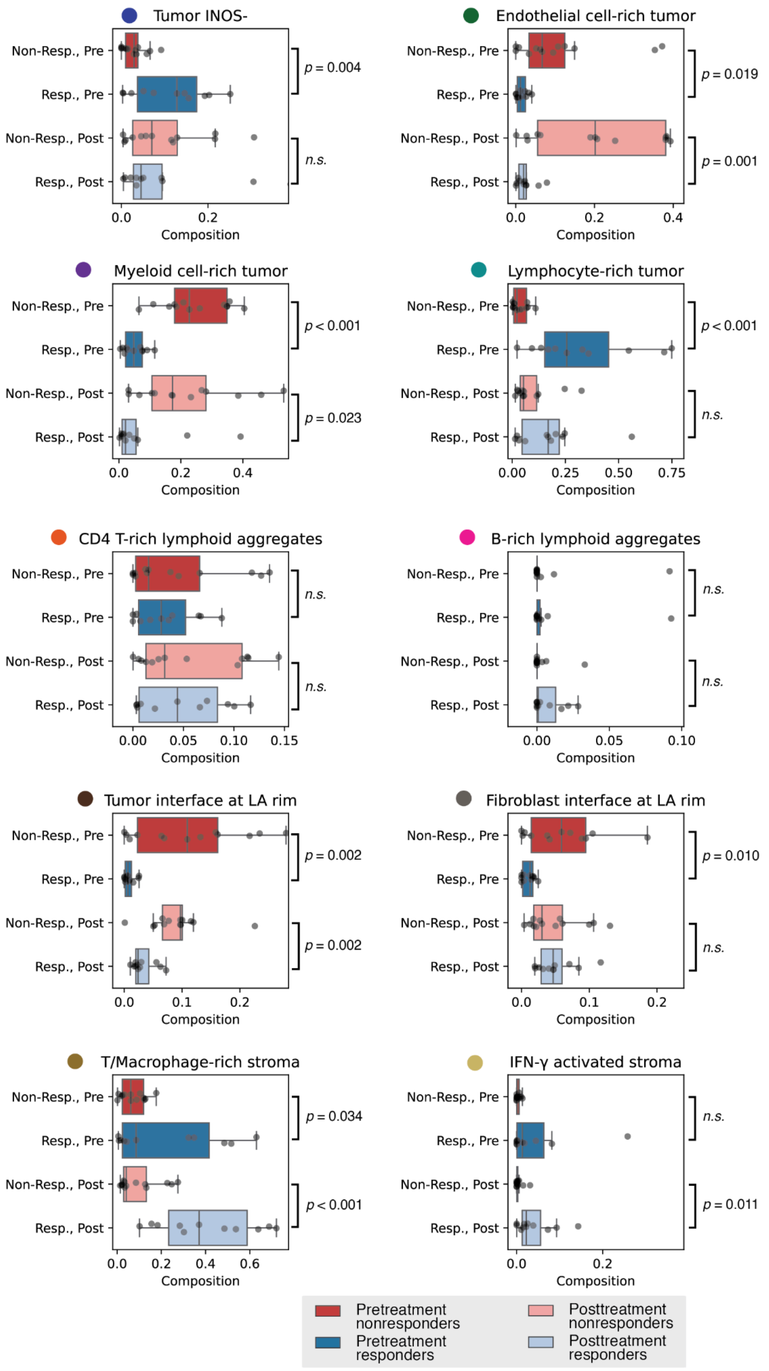


**Supplementary Fig. 8: Distinct enrichment patterns of microenvironment archetypes across patient groups**

The presence and frequency of major microenvironments are analyzed across conditions, including pretreatment vs. posttreatment and responders vs. nonresponders. Microenvironment archetypes exhibit distinct distributions across these conditions. Statistical significance is assessed using two-sample t-tests, and *p*-values are reported in the figure.

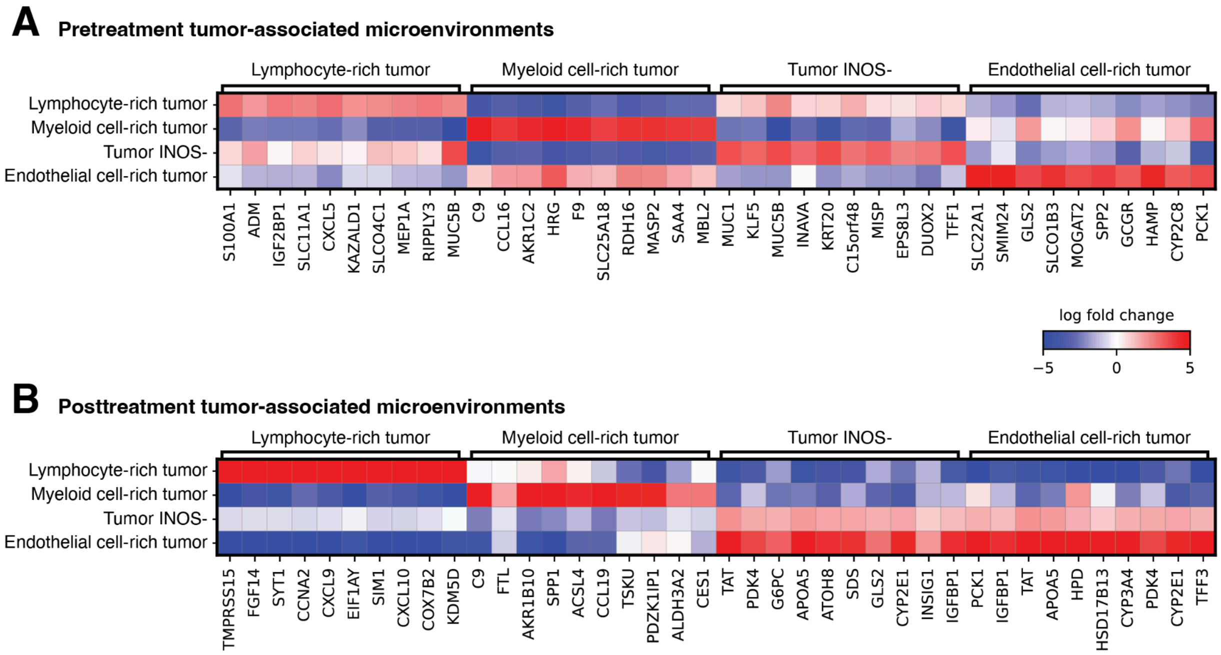


**Supplementary Fig. 9: Gene signatures of major tumor-associated microenvironments**

ST spots that map to predominantly one type of microenvironment during pseudo-spotting were extracted and compared against each other in a one-versus-rest analysis. Key genes for major tumor-associated microenvironments were identified in these comparisons for pretreatment and posttreatment samples. More targeted comparisons can be found in **Fig. 4** and **Fig. 5** of the main text.

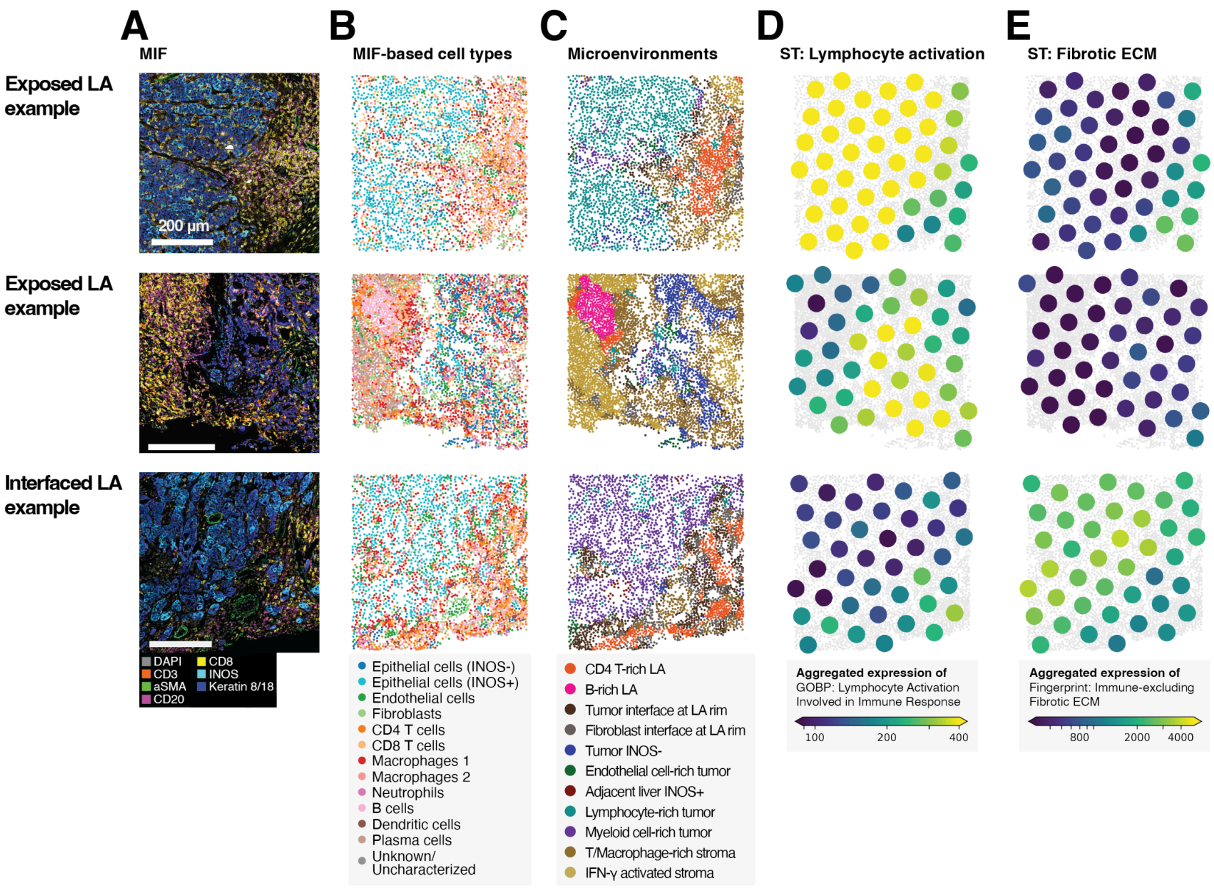


**Supplementary Fig. 10: Examples of LAs and their spatial contexts.**

The top two rows show exposed LAs and the bottom row shows an interfaced LA. **A.** MIF images showing strong stains of CD4, CD8, and CD20. **B.** Spatial plots of individual cells annotated by cell type. **C.** Spatial plots of individual cells annotated by microenvironment archetypes. Note the presence of LA-associated microenvironments. **D.** Spatial plots of ST spots colored by lymphocyte activation gene signatures. This signature is adapted from a GOBP term. **E.** Spatial plots of ST spots colored by fibrotic ECM gene signatures. This signature is curated from the fingerprint (**Methods**, **Supplementary Table 3**).

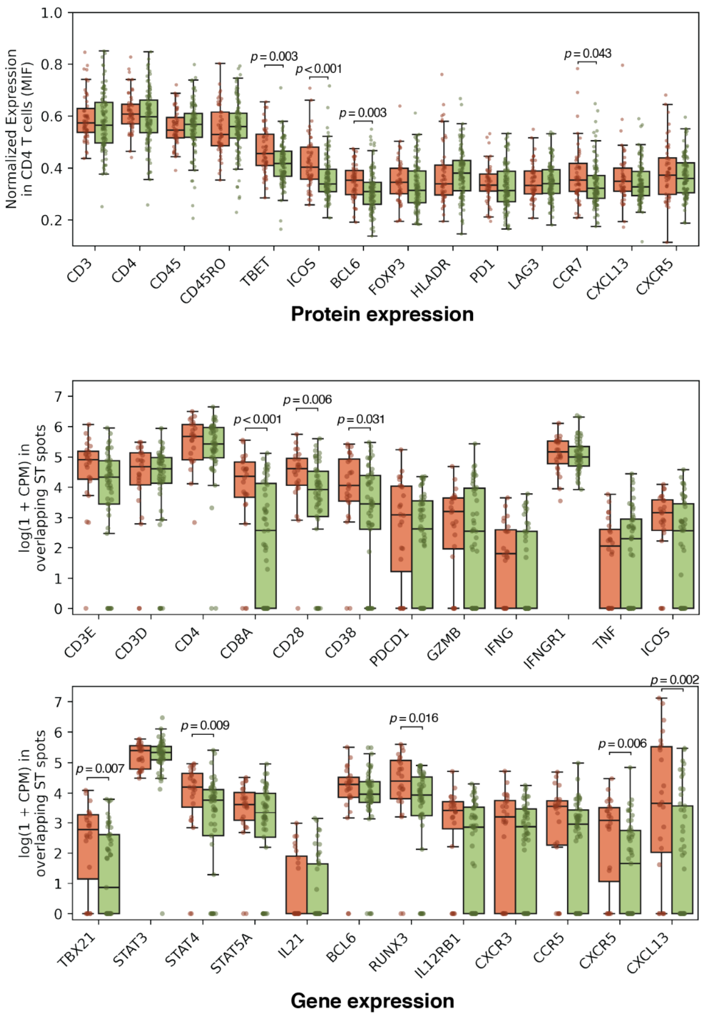


**Supplementary Fig. 11: Expression of key proteomic and transcriptomic biomarkers in exposed and interfaced LAs**

The box plots illustrate expression of key protein and gene biomarkers relevant to T cell functions in exposed (red) and interfaced (green) LAs. Protein biomarker expression was calculated in CD4 T cells identified from MIF data, while gene expression levels were calculated on ST spots mapped to LA-associated microenvironments.

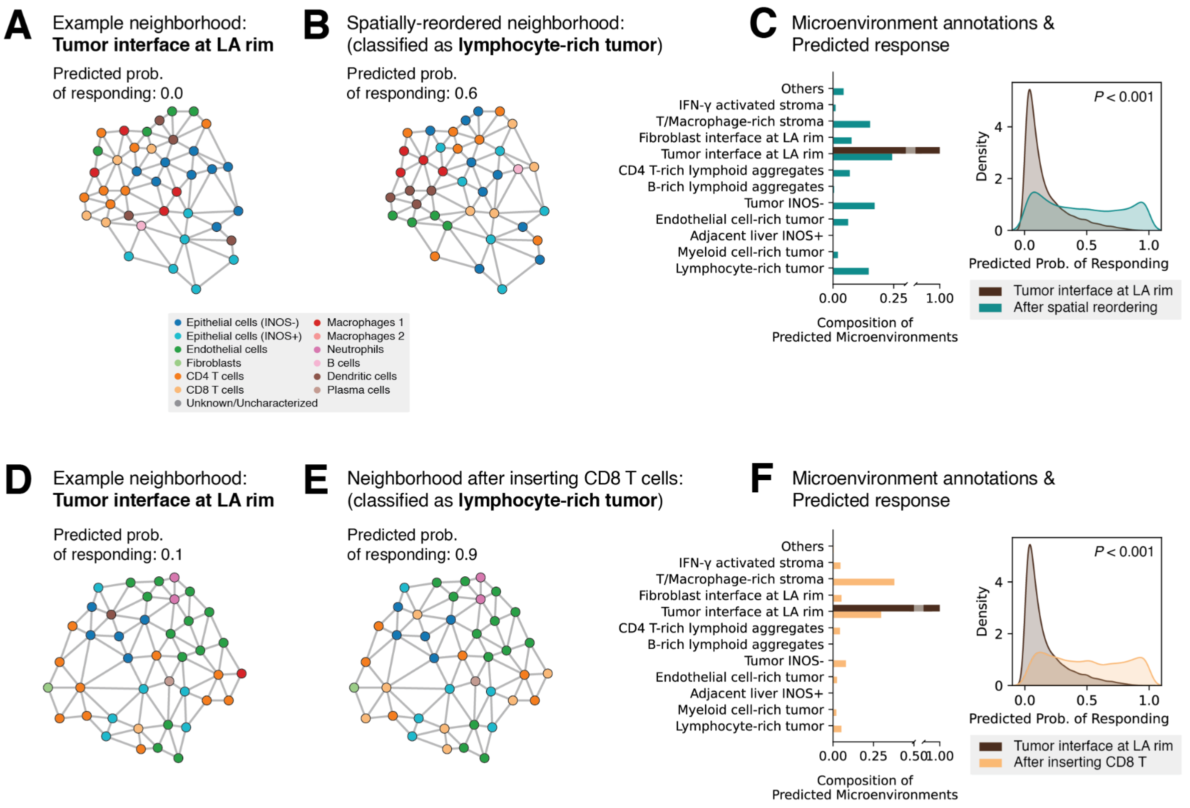


**Supplementary Fig. 12: Permutations of tumor interface at LA rims reveal resistance-associated spatial signatures**

**A-B**. In the first scenario, cells were spatially reordered to enforce mixing of tumor cells and lymphocytes. **C.** After permutation, more than two-thirds of the neighborhoods transitioned into either tumor-associated or stromal microenvironments (e.g., lymphocyte-rich tumor). Predicted probabilities of responding were significantly shifted to the right (Wilcoxon rank-sum test, $P < 0.001$). These findings suggest that lack of tumor-immune interactions is a defining feature of the tumor-LA interface structures and is strongly associated with nonresponse. **D-E**. In the second scenario, all immune cells from the neighborhood were extracted and randomly replaced with a target immune cell type, a process termed insertion. **F.** Insertion of CD8 T cells led to the transition from tumor interface microenvironments into response-indicative stromal microenvironments (i.e., fibroblasts with CD8 T & macrophages). Predicted probabilities were also significantly shifted to the right (Wilcoxon rank-sum test, $P < 0.001$), suggesting that CD8 T presence in the vicinity of LA is a key signature of exposed LAs.

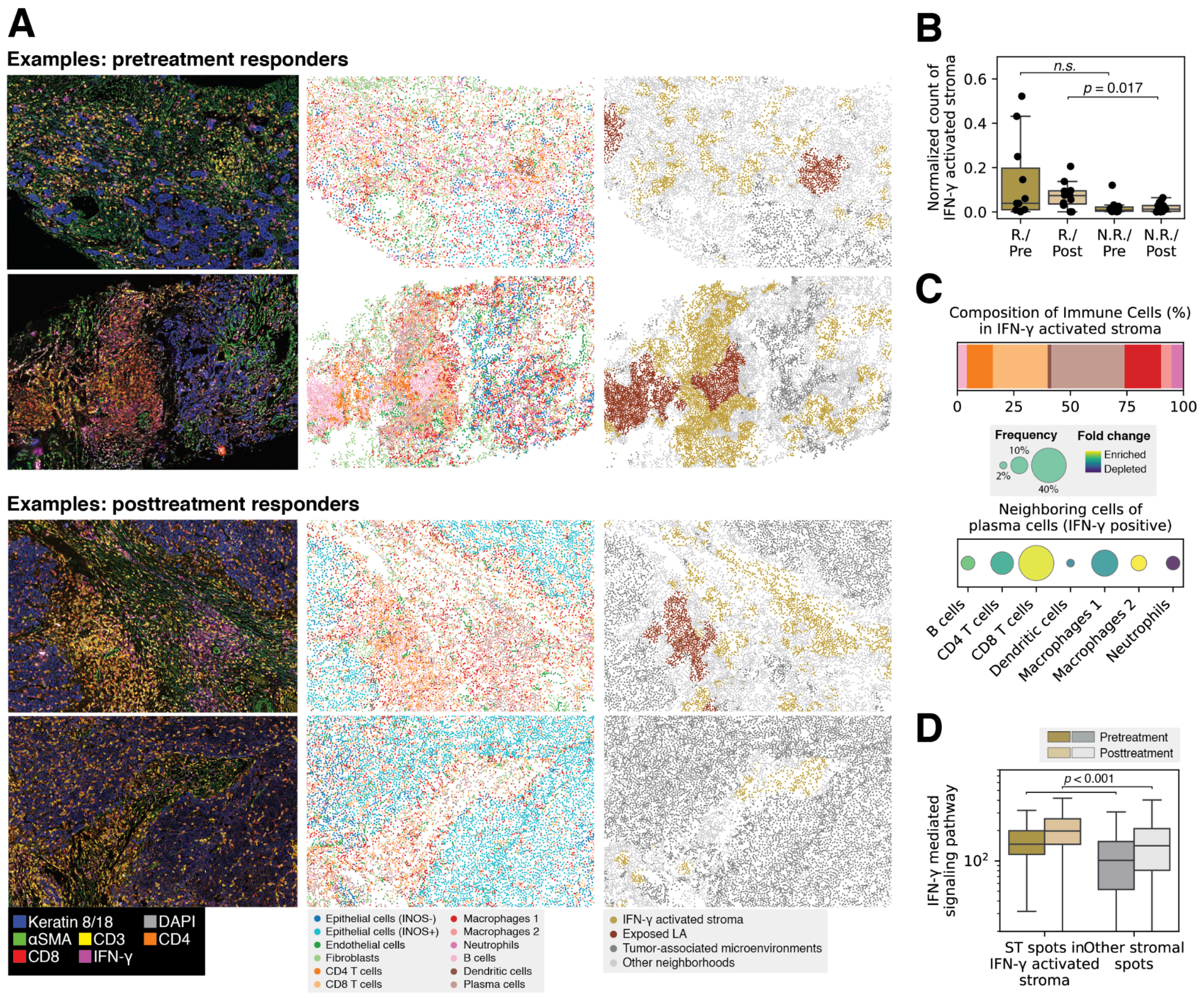


**Supplementary Fig. 13: Characterization of an IFN-γ-associated microenvironment archetype**

**A.** IFN-γ signals and IFN-γ associated microenvironments were observed in pretreatment and posttreatment responder samples. Regions from different conditions/patients are characterized and displayed as MIF images, cell type annotations, and microenvironment annotations. Note the presence of IFN-γ (magenta) in the MIF images, along with the co-localization of exposed LA and IFN-γ activated stroma microenvironments. **B.** The IFN-γ activated stroma microenvironment shows different enrichment across samples. Posttreatment responders have significantly more IFN-γ activated stroma than nonresponders. **C.** Composition and enrichment of immune cells in IFN-γ associated stroma microenvironments. Observations on IFN-γ positive plasma cells show enriched interactions with CD8 T cells. **D.** ST spots spatially overlapped with IFN-γ activated stroma microenvironments also exhibit elevated gene expression levels in the associated signaling pathway. An overall increase in IFN-γ signaling activities is observed following treatment in the stroma compartments of both responders and non-responders.

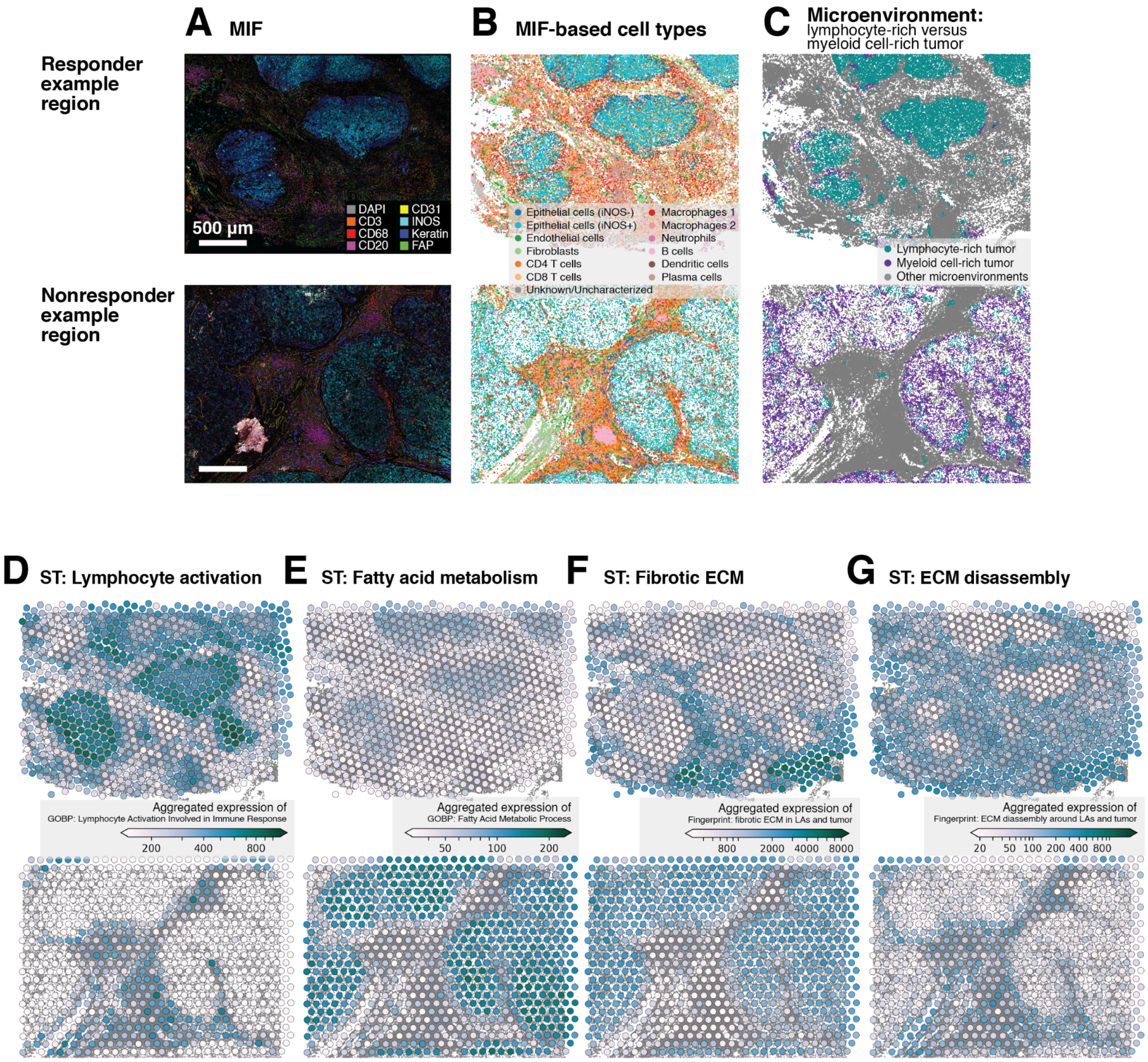


**Supplementary Fig. 14: Characterizations of two INOS-positive tumor regions**

**A.** MIF images. **B.** Spatial plots of individual cells annotated by cell type. **C.** Spatial plots of individual cells annotated by microenvironment archetypes. Note the contrast of lymphocyte-rich tumor and myeloid cell-rich tumor. **D.** ST spots colored by lymphocyte activation gene signatures, adapted from a GOBP term. **E.** ST spots colored by fatty acid metabolism gene signatures, adapted from a GOBP term. **F.** ST spots colored by fibrotic ECM gene signatures. This signature is curated from the fingerprint (**Methods**, **Supplementary Table 3**). **G.** ST spots colored by ECM disassembly gene signatures. This signature is curated from the fingerprint.

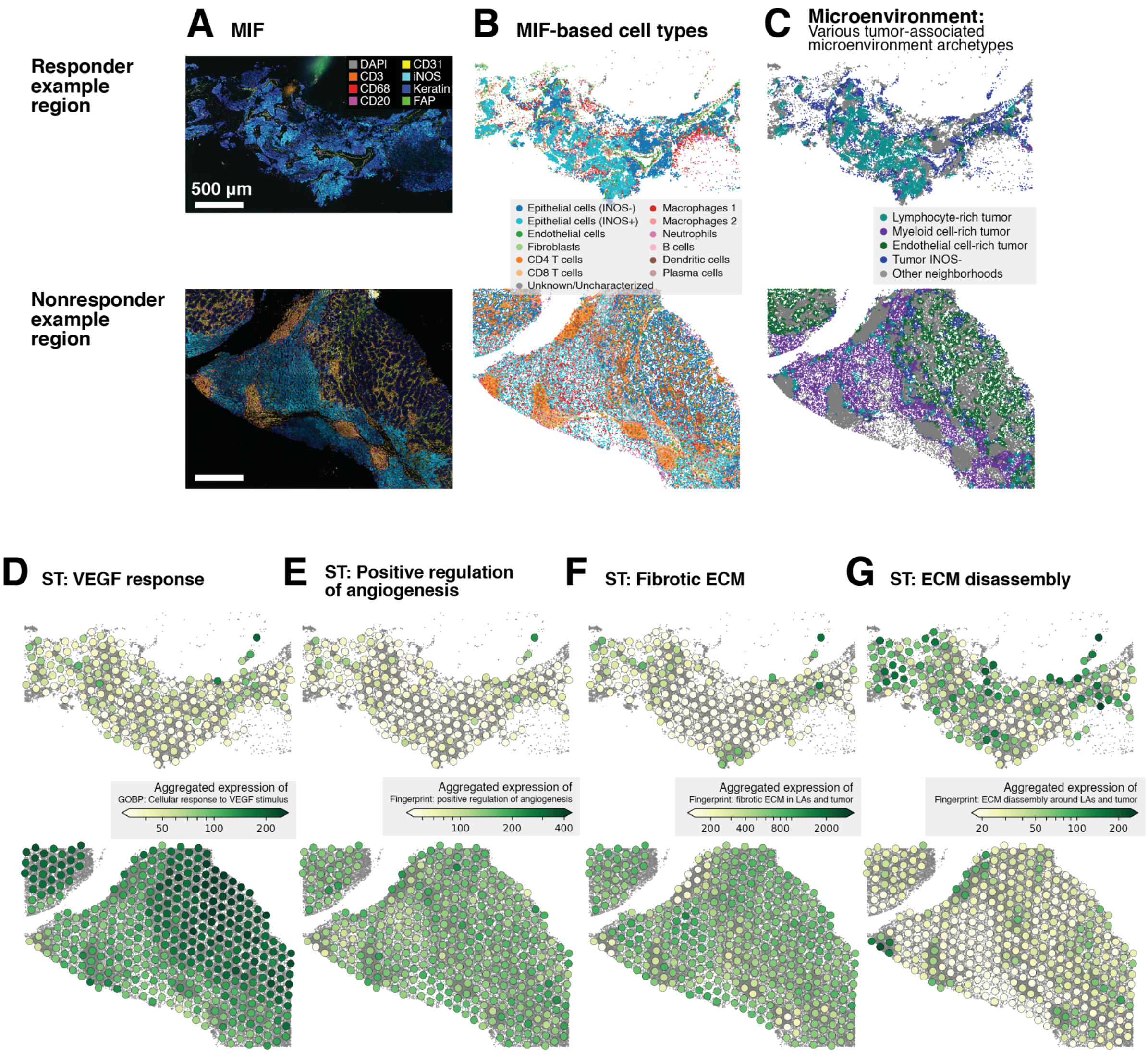


**Supplementary Fig. 15: Characterizations of two INOS-negative tumor regions**

**A.** MIF images. **B.** Spatial plots of individual cells annotated by cell type. **C.** Spatial plots of individual cells annotated by microenvironment archetypes. Note the presence of different tumor-associated microenvironments. **D.** ST spots colored by cellular response to VEGF gene signatures, adapted from a GOBP term. **E.** ST spots colored by angiogenesis regulation gene signatures. This signature is curated from the fingerprint (**Methods**, **Supplementary Table 3**). **F.** ST spots colored by fibrotic ECM gene signatures. This signature is curated from the fingerprint. **G.** ST spots colored by ECM disassembly gene signatures. This signature is curated from the fingerprint.

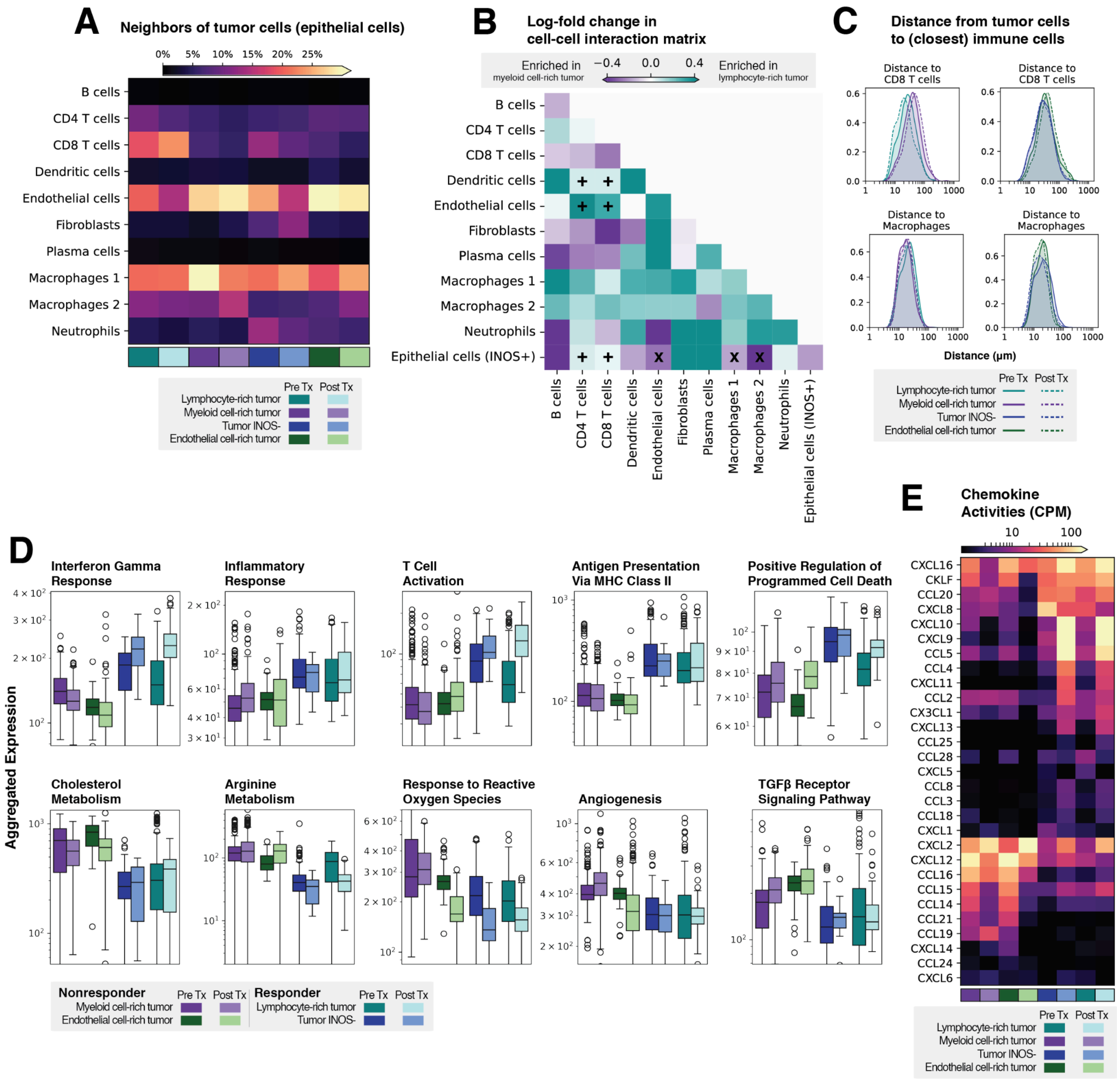


**Supplementary Fig. 16: Detailed cellular and transcriptomic features of tumor-associated microenvironments**

**A.** Neighboring cells of tumor cells (i.e., epithelial cells) in all tumor-associated microenvironments before and after immunotherapy treatment. Lymphocyte-rich tumor microenvironments exhibit more CD8 T-tumor interactions, while myeloid cell-rich and endothelial cell-rich tumor microenvironments have more macrophage-tumor and endothelium-tumor interactions. **B.** Differential enrichment of cell-cell interactions between lymphocyte-rich and myeloid cell-rich tumor microenvironments. Each entry represents the log-fold change in the frequency of neighboring pairs of cells, normalized by average compositions of cell types in the two microenvironments. Lymphocyte-rich tumor microenvironments exhibit more CD4 T-tumor, CD8 T-tumor, T-DC and T-endothelium interactions (plus marks). Myeloid cell-rich tumor microenvironments, on the contrary, have more macrophage-tumor and endothelium-tumor interactions (cross marks). **C.** Distributions of tumor-immune distances before and after immunotherapy treatment. Immune environments of tumor cells are characterized by their closest distances to different types of immune cells. Lymphocyte-rich tumor microenvironments see closer distances between tumor cells and CD8 T cells, which decrease further after treatment. **D.** Transcriptomic states of tumor-associated microenvironments characterized by a comprehensive set of gene signatures. Stronger immune activities were observed in lymphocyte-rich tumor and tumor INOS- microenvironments, especially following treatment, as exemplified by the elevated signatures associated with IFN-γ responses, T cell activation, antigen presentation, and inflammation. On the contrary, myeloid cell-rich and endothelial cell-rich tumor microenvironments have strong signatures of metabolism(i.e., fatty acid, cholesterol, and arginine metabolism), reactive oxygen species, angiogenesis, and TGFβ signaling. **E.** Expression of chemokines in tumor-associated microenvironments. Lymphocyte-rich tumor microenvironments have high expression of *CXCL16*, *CXCL10*, *CXCL13*, *CXCL9*, and *CCL5*, most of which are known recruiters of T cells. Myeloid cell-rich tumor shows elevated signals of *CXCL2*, *CXCL12*, and *CCL16*, which attract immune cells including neutrophils and monocytes.

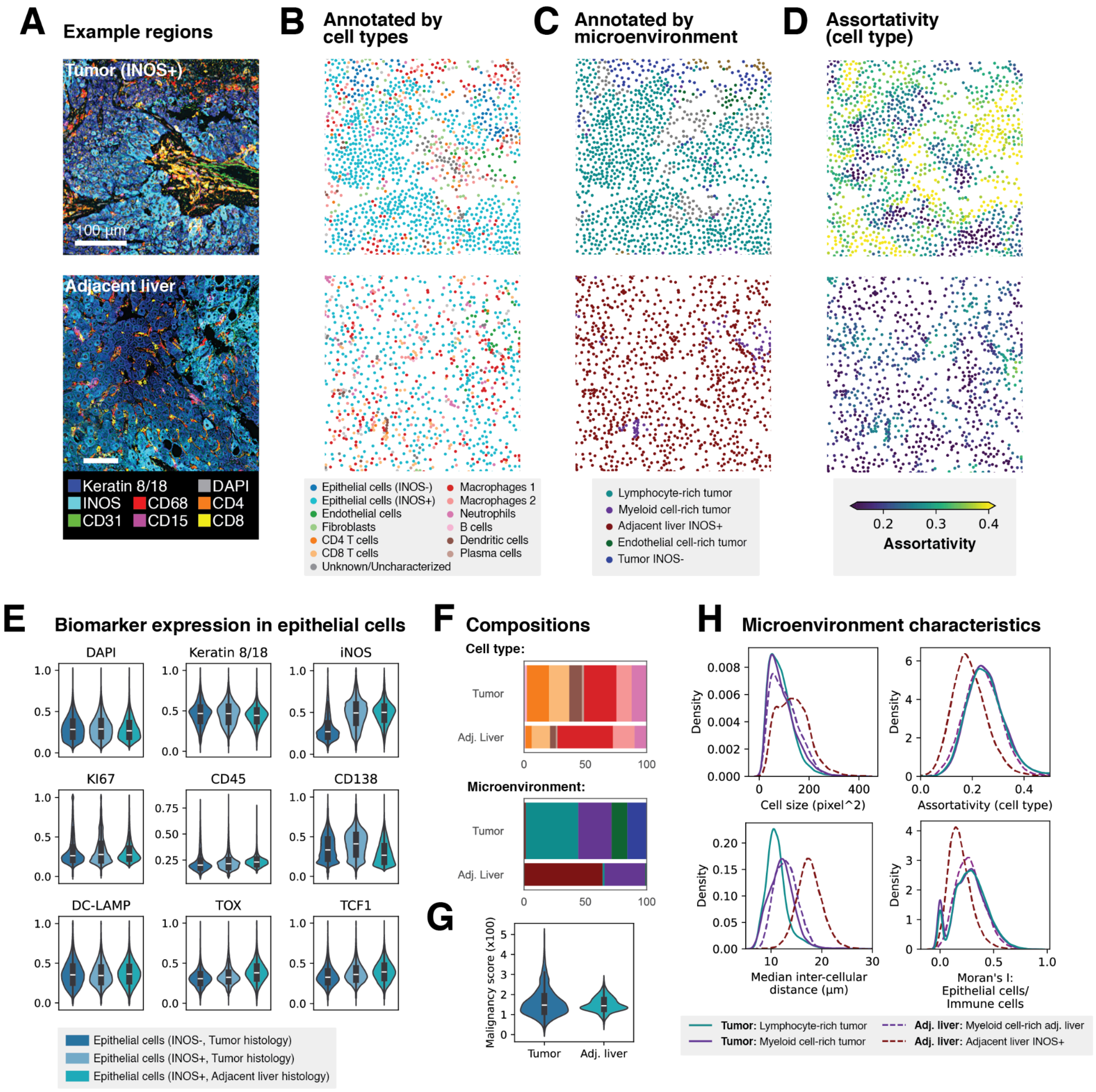


**Supplementary Fig. 17: Distinct cell types and microenvironment characteristics observed across different histology compartments**

**A.** Representative regions from tumor and tumor-adjacent liver compartments. Note that both regions have high expression of INOS. **B.** Spatial plots of cells in the regions colored by cell types. The tumor-adjacent liver compartment contains more dispersed macrophages (resident macrophages). **C.** Spatial plots of cells in the regions colored by microenvironments. Tumor and tumor-adjacent liver compartments have distinct compartment-specific microenvironments. **D.** Spatial plots of cells in the regions colored by local assortativities. Assortativity is a metric measuring the coherency of cell types in the neighborhood of the corresponding cell. High assortativities indicate neighborhoods composed of higher proportions of a single type of cells. **E.** Protein expression profiles of epithelial cells from different histology compartments. All epithelial cells have high expression of keratin 8/18. Both tumor-adjacent liver and tumor compartments contain epithelial cells with strong signals of INOS, though no biomarkers clearly distinguish tumoral and tumor-adjacent liver epithelial cells. **F.** Compositions of cell types and microenvironments in different histology compartments. Note the presence of compartment-specific microenvironments (cyan, red, green, blue). All analyses in the main texts regarding myeloid cell-rich tumor microenvironment (purple) are performed on neighborhoods of tumor histology. **G.** Distributions of malignancy scores from the two compartments are similar and show no significant differences. **H.** Distinct characteristics of compartment-specific microenvironments. Tumor-specific and tumor-adjacent liver-specific microenvironments show differences in cell sizes, inter-cellular distances, assortativities of cell types as plotted in panel **D**, and spatial autocorrelation.

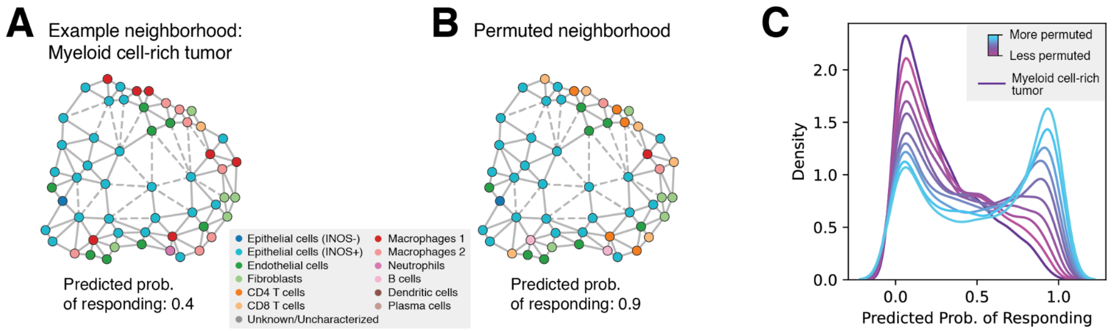


**Supplementary Fig. 18: Permutations of the immune environment in myeloid cell-rich tumor microenvironments reduce resistance signals**

**A.** An example neighborhood of myeloid cell-rich tumor microenvironment plotted as a graph. **B.** Computational permutations performed on the neighborhood altered its immune environment by replacing myeloid cells (macrophages and neutrophils) with lymphocytes (B cells and T cells). This permutation increased the predicted probability of response to immunotherapy to 0.9. **C.** Distributions of predicted probabilities of response before and after permutations. We applied the same permutation to a larger set of myeloid cell-rich tumor microenvironments randomly sampled from the data. Notably, introducing more lymphocytes into the neighborhoods progressively shifts the prediction distributions to the right, indicating an increased likelihood of response.

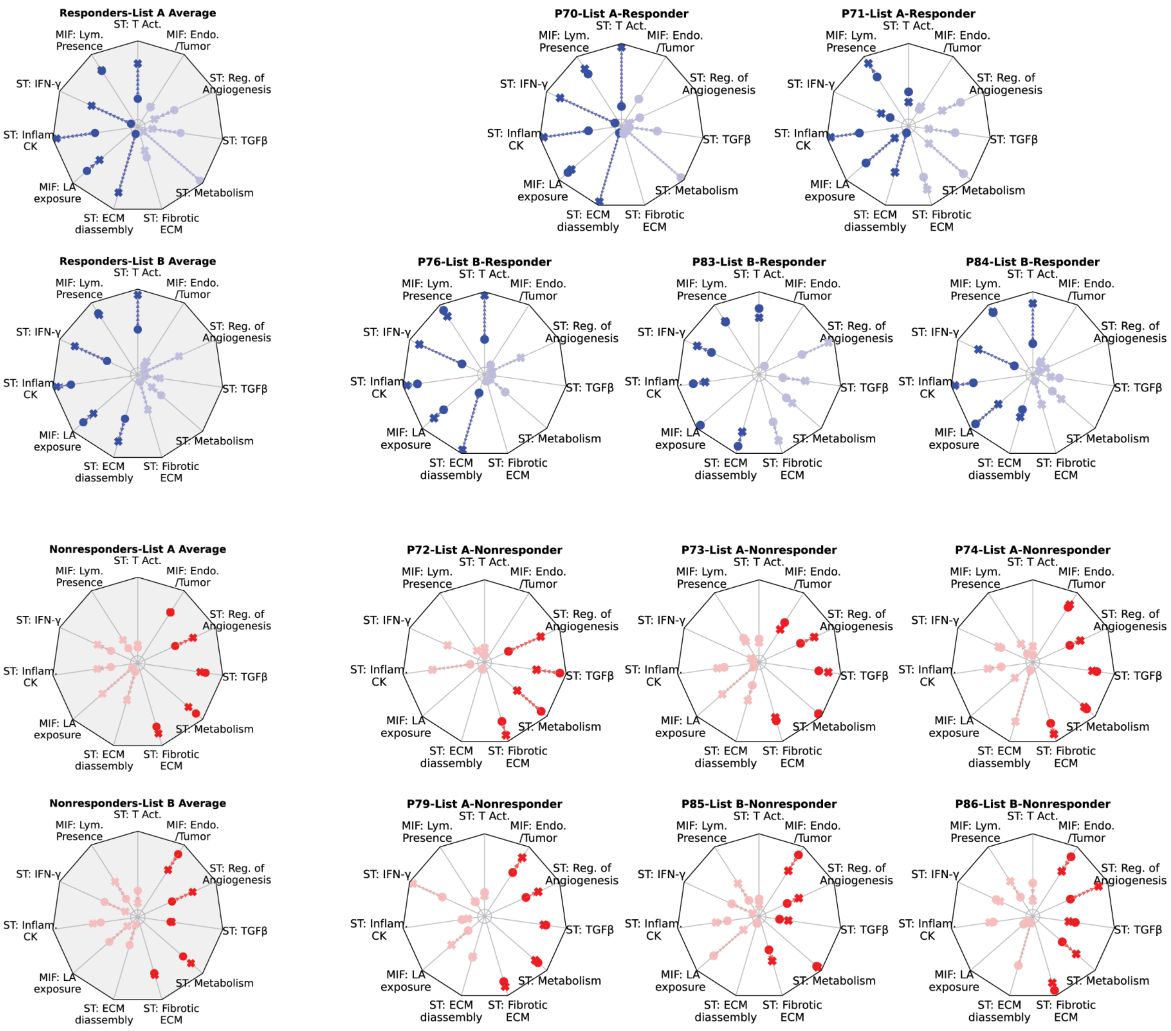


**Signatures (counterclockwise):**
ST: T Act. - *Intratumoral T cell activation*
MIF: Lym.Presence - *Intratumoral lymphocyte presence*
ST: IFN-γ - *Response to IFN-γ in LAs and lymphocyte-rich tumor*
ST: Inflam. CK - *Inflammatory chemokines*
MIF: LA exposure - *Lymphoid aggregate exposure*
ST: ECM disassembly -*ECM disassembly and remodeling around LAs and tumor*
ST: Fibrotic ECM - *Immune-excluding fibrotic ECM in LAs and tumor*
ST: Metabolism - *Metabolism programs*
ST: TGF-β - *TGFβ signaling and response*
ST: Reg. of Angiogenesis - *Regulation of angiogenesis in tumor*
MIF: Endo./Tumor - *Tumor-endothelium interaction*

**Supplementary Fig. 19: Heterogeneous fingerprints of response and resistance across patients**

We calculated the multi-omic fingerprint defined based on microenvironment analysis (**Fig. 6**, **Methods**) on MIF and ST data of each patient. Averages over responders and nonresponders receiving list A (anti-PD1) or list B (anti-PD1 + anti-CTLA4) treatment were calculated and displayed in the leftmost column. Responders consistently showed elevated IFN-γ and inflammatory chemokine signatures posttreatment, coupled with high levels of intratumoral T cell activities, LA exposure, and ECM disassembly signals. In contrast, nonresponders displayed reduced T cell activity and were characterized by resistance signatures, including immune-excluding fibrotic ECM, angiogenic activity, TGFβ signaling, and enhanced metabolism. These resistance signatures were also highly heterogeneous across individual patients.

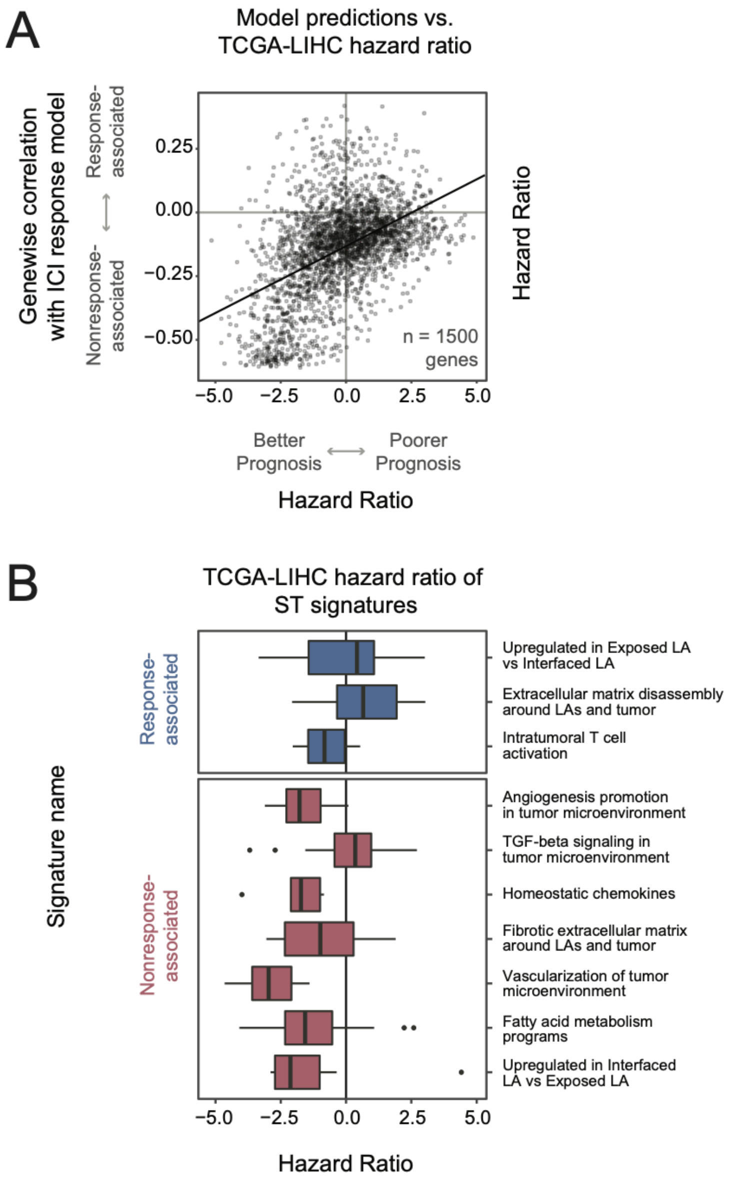


**Supplementary Fig. 20: Nonresponse-associated ST signatures are associated with improved overall survival outside of the immunotherapy setting.**

**A.** The overall relationship between response association between TCGA liver hepatocellular carcinoma (TCGA-LIHC) hazard ratio and gene-level immune checkpoint inhibition (ICI) prediction correlations from ST-MLP. Genes with high correlations are more associated with response, and vice versa. Many genes associated with ICI nonresponse predict better prognosis in TCGA-LIHC. **B.** TCGA-LIHC hazard ratio of ST signatures identified in this study. Several nonresponse signatures have a bias towards better prognosis, while response signatures are more neutral.

**Supplementary Table 1: Summary of spatially co-registered multi-omics data of HCC responders and nonresponders**

| Patient | Response | ST data | MIF data |
|---|---|---|---|
| P70 | Responder | Pre Tx: **Avail.**<br>Post Tx: **Avail.** | Pre Tx: **2** FOV<br>Post Tx: **2** FOV |
| P71 | Responder | Pre Tx: **Avail.**<br>Post Tx: **Avail.** | Pre Tx: **2** FOV, **2** FOV aligned to ST<br>Post Tx: **2** FOV |
| P72 | Nonresponder | Pre Tx: **Avail.**<br>Post Tx: **Avail.** | Pre Tx: **2** FOV, **2** FOV aligned to ST<br>Post Tx: **1** FOV, **1** FOV aligned to ST |
| P73 | Nonresponder | Pre Tx: **Avail.**<br>Post Tx: **Avail.** | Pre Tx: **2** FOV, **2** FOV aligned to ST<br>Post Tx: **2** FOV, **1** FOV aligned to ST |
| P74 | Nonresponder | Pre Tx: **Avail.**<br>Post Tx: **Avail.** | Pre Tx: **2** FOV, **2** FOV aligned to ST<br>Post Tx: **2** FOV, **2** FOV aligned to ST |
| P75 | Responder | Pre Tx: **N/A**<br>Post Tx: **N/A** | Pre Tx: **2** FOV<br>Post Tx: **1** FOV |
| P76 | Responder | Pre Tx: **Avail.**<br>Post Tx: **Avail.** | Pre Tx: **2** FOV, **2** FOV aligned to ST<br>Post Tx: **2** FOV, **2** FOV aligned to ST |
| P77 | Nonresponder | Pre Tx: **N/A**<br>Post Tx: **N/A** | Pre Tx: **2** FOV<br>Post Tx: **2** FOV |
| P79 | Nonresponder | Pre Tx: **Avail.**<br>Post Tx: **Avail.** | Pre Tx: **2** FOV, **1** FOV aligned to ST<br>Post Tx: **2** FOV, **2** FOV aligned to ST |
| P83 | Responder | Pre Tx: **Avail.**<br>Post Tx: **Avail.** | Pre Tx: **1** FOV, **1** FOV aligned to ST<br>Post Tx: **2** FOV |
| P84 | Responder | Pre Tx: **Avail.**<br>Post Tx: **Avail.** | Pre Tx: **2** FOV, **2** FOV aligned to ST<br>Post Tx: **2** FOV, **2** FOV aligned to ST |
| P85 | Nonresponder | Pre Tx: **Avail.**<br>Post Tx: **Avail.** | Pre Tx: **1** FOV, **1** FOV aligned to ST<br>Post Tx: **2** FOV |
| P86 | Nonresponder | Pre Tx: **Avail.**<br>Post Tx: **Avail.** | Pre Tx: **2** FOV, **2** FOV aligned to ST<br>Post Tx: **2** FOV |
| Total:<br>**13** patients | **6** responders<br>**7** nonresponders | **11** Pre Tx<br>**11** Post Tx | **24** Pre Tx FOV, **17** FOV aligned to ST<br>**24** Post Tx FOV, **10** FOV aligned to ST |

**Supplementary Table 2: Antibodies used in the 51-plex PhenoCycler imaging**

| Antibody | Vendor | Reporter | Antibody | Vendor | Reporter |
|---|---|---|---|---|---|
| αSMA | R&D | AF488 | CXCL13 | Abcam | Cy5 |
| Bcl-6 | Lifespan | Cy5 | CXCR5 | CST-Custom | Atto550 |
| CCR7 | Abcam | AF488 | DNA | - | DAPI |
| CD107a | Akoya | Cy5 | DC-LAMP | Novus Biologicals | AF488 |
| CD11c | Akoya | Cy5 | EOMES | eBioscience | Atto550 |
| CD138 | Abcam | Atto550 | FAP | R&D | AF488 |
| CD14 | Abcam | AF488 | FoxP3 | Thermofisher | AF488 |
| CD141 | Akoya | Atto550 | Gr-B | Akoya | Atto550 |
| CD15 | BioLegend | Cy5 | HLA-DR | Abcam | AF488 |
| CD163 | Novus Biologicals | Atto550 | ICOS | CST | Cy5 |
| CD1c | Akoya-Abcam | AF647 | IDO1 | Akoya | Cy5 |
| CD20 | Akoya | AF488 | IFN-g | Akoya | Atto550 |
| CD204 | Amsbio | AF488 | iNOS | Akoya | Atto550 |
| CD206 | Lifespan | Atto550 | Keratin 8/18 | Akoya | AF647 |
| CD209 | Akoya-Abcam | Atto550 | Ki67 | Akoya | Atto550 |
| CD21 | Akoya | Atto550 | Lag3 | Lifespan | Cy5 |
| CD3 | Dako | AF647 | PD-1 | Leinco | AF488 |
| CD31 | Akoya | AF488 | PD-L1 | Abcam | Atto550 |
| CD34 | Abcam | AF488 | PNad | BioLegend | AF488 |
| CD4 | Akoya | Cy5 | Podoplanin | Akoya-Abcam | AF647 |
| CD45 | Akoya | Cy5 | T-bet | BioLegend | AF488 |
| CD45RO | Akoya | Atto550 | TCF1 | Akoya | AF647 |
| CD56 | Leinco | Cy5 | TOX | Akoya | Atto550 |
| CD68 | Akoya | Cy5 | VISTA | Akoya | Atto550 |
| CD79a | Akoya | AF647 | XCR1 | CST | Atto550 |
| CD8 | Akoya | Atto550 | | | |

**Supplementary Table 3: Spatial transcriptomics signatures for immunotherapy resistance**

| Name | Description | Content |
|---|---|---|
| ST: T Act. | Intratumoral T cell activation | GOBP terms: Antigen Processing And Presentation Of Peptide Antigen Via MHC Class II (GO:0002495) and T Cell Activation Involved In Immune Response (GO:0002286) |
| ST: IFN-γ | Response to IFN-γ/Type II interferon | GOBP terms: Cellular Response To Type II Interferon (GO:0071346) and Type II Interferon-Mediated Signaling Pathway (GO:0060333) |
| ST: Inflam. CK | Inflammatory chemokines | Difference between expression of: (inflammatory set) CXCL16, CXCL9, CCL5, CXCL10, CCL4, CXCL13, CCL18, CXCL11 and (homeostatic set) CCL16, CCL14, CCL15, CCL19, CCL21, CXCL14 |
| ST: ECM disassembly | ECM disassembly and remodeling | ADAMTS1,ADAMTS4,ADAMTS10,ADAMTS20,ADAMTS8,CAPG,CCDC80,CTSV,LOXL1,LOXL4,MMP1,MMP10,MMP11,MMP12,MMP14,MMP2,MMP7,MMP9 |
| ST: Fibrotic ECM | Immune-excluding ECM | BGN,COL14A1,COL1A1,COL1A2,COL3A1,COL4A1,COL4A2,COL5A1,COL6A1,COL6A2,COL6A3,DCN,DPT,ELN,FBLN5,FGA,FGB,FGG,HAS2,LAMA2,LAMC3,LUM,PLG,SERPINE1,SPARC,THBS1,THBS2,TTR |
| ST: Metabolism | Combinations of metabolism programs | ABAT,ABCB11,ABCD3,ACAA2,ACACA,ACADSB,ACOX2,ACSL1,ACSL3,ACSL4,ACSM2A,ACSM2B,ACSM3,ADIPOR2,AKR1C2,AKR1C4,AKR1D1,AMACR,ANGPTL3,APOA1,APOA2,APOF,ARG1,ASS1,BDH1,CBR1,CPT1A,CYP2A6,CYP2B6,CYP2C9,CYP2E1,CYP2J2,CYP3A7,CYP4A11,CYP4F11,CYP4F2,CYP51A1,CYP7A1,CYP7B1,CYP8B1,DDAH1,DHCR7,ECHDC3,FASN,FDFT1,GATM,GGT5,GLYAT,GPAM,HAO2,HMGCR,HMGCS1,HPGD,HSD11B1,HSD17B14,IGF1,INSIG1,KYNU,LIPC,MSMO1,MVK,NPC1L1,NSDHL,PCCB,PRLR,RDH16,SLC27A2,SLC27A5,SQLE,SRD5A1,SULT2A1,TM7SF2,TSKU,VNN1 |
| ST: TGFβ | TGFβ signaling and response | ANKRD1,APOA1,BAMBI,COL1A2,COL3A1,ENG,FMOD,FOS,GDF15,LEFTY1,LRRC32,LTBP2,SOX6,TGFB2,WNT4,WNT7A |
| ST: Interface | Upregulated in Interface LA | ALB, SPP1, CES1, SERPINA1, SERPINA3, FGA, FGB, FGG, APOA1, APOA2, ORM1, GSTA2, VTN, UGT2B4, C3 |
| ST: Reg. of Angiogenesis | Positive angiogenesis regulation | EMILIN1,RAMP2,STAB1,AQP1,RHOB,NAXE,CLDN5,HMOX1,HYAL1,VASH1,MYDGF,HSPG2,EPHA1,ERBB2,HTATIP2,VEGFC,ECSCR,CDH5,ANGPTL3,DCN,ADGRA2,EMP2,THBS4,KDR,FLT1,NPR1,APLNR |
